# ASH1L REGULATES THE STRUCTURAL DEVELOPMENT OF NEURONAL CIRCUITRY BY MODULATING BDNF/TrkB SIGNALING IN HUMAN NEURONS

**DOI:** 10.1101/2020.02.18.954586

**Authors:** Seon Hye Cheon, Allison M. Culver, Anna M. Bagnell, Foster D. Ritchie, Janay M. Clytus, Mikayla McCord, Carin M. Papendorp, Evelyn Chukwurah, Austin J. Smith, Mara H. Cowen, Pankaj S. Ghate, Shannon W. Davis, Judy S. Liu, Sofia B. Lizarraga

**Author notes:** These authors contributed equally to this work (Allison M. Culver, Anna M. Bagnell). Corresponding author and Lead contact Sofia B. Lizarraga, PhD.

## Abstract

Autism spectrum disorders (ASD) are associated with defects in neuronal connectivity and are highly heritable. Genetic findings suggest that there is an overrepresentation of chromatin regulatory genes among the genes associated with ASD. ASH1 like histone lysine methyltransferase (ASH1L) was identified as a major risk factor for autism. ASH1L methylates Histone H3 on Lysine 36, which is proposed to result primarily in transcriptional activation. However, how mutations in ASH1L lead to deficits in neuronal connectivity associated with autism pathogenesis is not known. We report that ASH1L regulates neuronal morphogenesis by counteracting the catalytic activity of Polycomb Repressive complex 2 group (PRC2) in stem cell-derived human neurons. Depletion of ASH1L decreases neurite outgrowth and decreases expression of the gene encoding the neurotrophin receptor TrkB whose signaling pathway is linked to neuronal morphogenesis. This is overcome by inhibition of PRC2 activity, indicating a balance between the Trithorax group protein ASH1L and PRC2 activity determines neuronal morphology and connectivity. Thus, ASH1L epigenetically regulates neuronal connectivity by modulating the BDNF-TrkB signaling pathway, which likely contributes to the neurodevelopmental pathogenesis associated with ASD in patients with ASH1L mutations.

**eTOC BLURB:** Cheon et al. report a novel epigenetic mechanism that implicates the counteracting activities of the evolutionarily conserved Trithorax (ASH1L) and Polycomb (PRC2) chromatin regulators, in the modulation of human neuronal connectivity by regulating the developmentally important TrkB-BDNF signaling pathway.

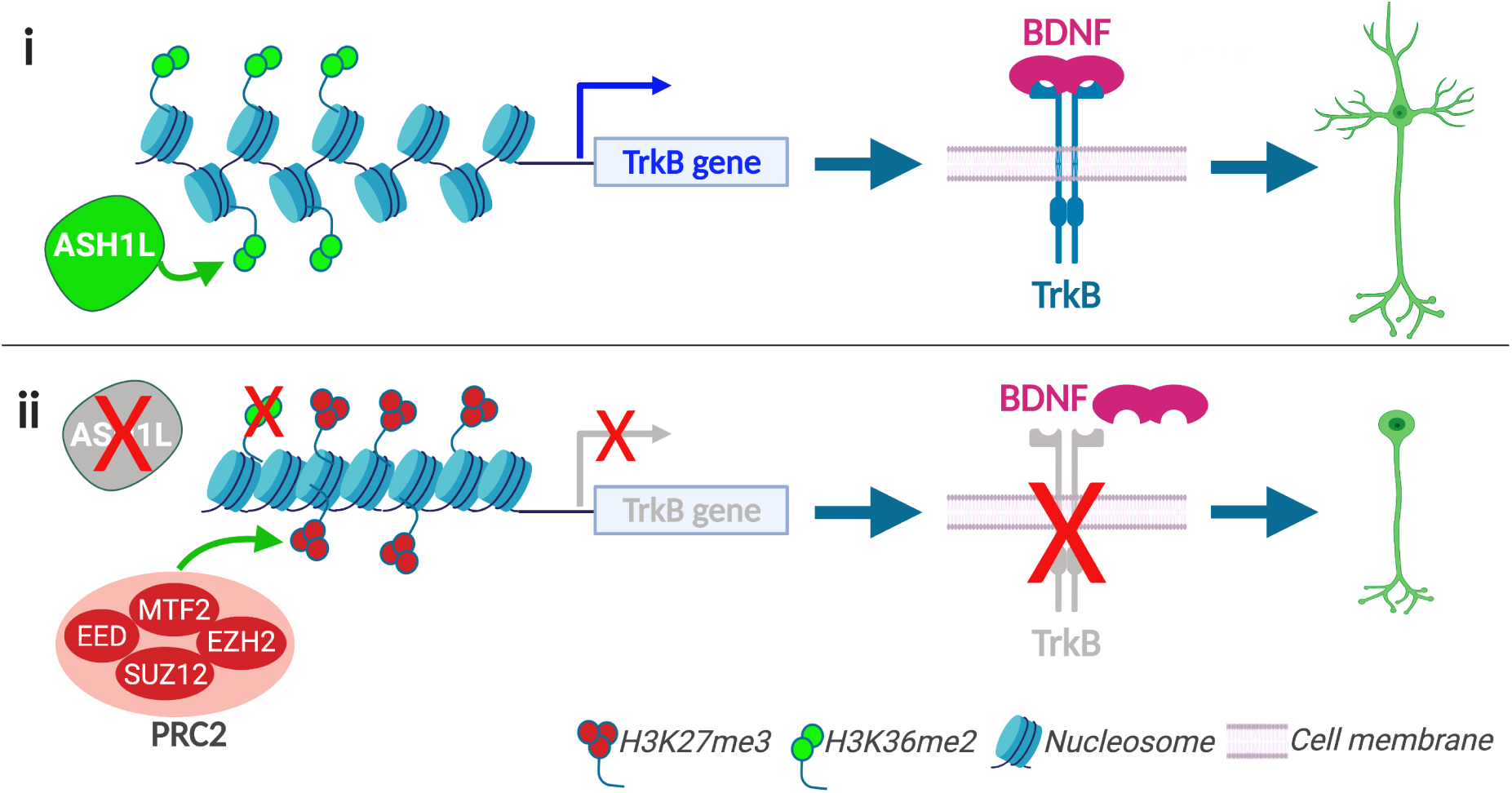

**HIGHLIGHTS:**

- ASH1L regulates neuronal morphogenesis by modulating neurotrophin signaling
- Counteracting activities of Trithorax (ASH1L) and Polycomb (PRC2) affect neuronal arborization
- Loss of ASH1L modulates growth cone size in human neurons

## INTRODUCTION

Autism spectrum disorder (ASD) affects 1 in 59 children and represents a growing public health concern worldwide. Between 50 to 75% of ASD cases are of unknown or complex genetic etiology. Thus suggesting a role for epigenetic mechanisms underlying the pathogenesis of ASD (Kuehner et al., 2019). Epigenetic mechanisms entail the incorporation or removal of chemical modifications on histones or genomic DNA that lead to the repression or activation of genes. Mutations in epigenetic regulators, including histone and DNA methylators, have been associated with ASD (Lamonica and Zhou, 2019) (Kuehner et al., 2019). In particular, there is a significant over-representation of chromatin regulators (over 40%) among genes that contain high risk variants for ASD, according to the SFARI curated database of autism genes (Abrahams et al., 2013). In particular, de novo truncating and missense mutations in ASH1L have been identified in large autism cohorts (De Rubeis et al., 2014; Tammimies et al., 2015; Wang et al., 2016). Furthermore, recent evidence suggests that ASH1L is associated with severe forms of autism, as patients with mutations in ASH1L suffer from intellectual disabilities, speech difficulties, seizures and postnatal microcephaly (failure of the brain to grow postnatally) (Faundes et al., 2017; Okamoto et al., 2017; Stessman et al., 2017). Therefore, the human phenotypes associated with mutations in ASH1L suggest a major role for this chromatin modifier during the development of neuronal connectivity.

ASH1L is a member of the Trithorax group of proteins. Trithorax proteins are major regulators of embryonic development that are proposed to antagonize the activity of the Polycomb repressor complex (Schuettengruber et al., 2017). In particular, ASH1L dimethylation of Histone H3 in lysine 36 (H3K36me2) antagonizes the activity of the Polycomb Repressor Complex 2 (PRC2) by preventing the trimethylation of Histone H3 Lysine 27 (H3K27me3) (Huang and Zhu, 2018; Miyazaki et al., 2013). The function of PRC2 in neuronal development has been associated with cell fate control of neurogenesis (Corley and Kroll, 2015; Pereira et al., 2010), and more recently, has been linked to neuronal maturation, as its catalytic subunit Enhancer of Zeste Homolog 2 (EZH2) is proposed to modulate neuronal arborization (Qi et al., 2014). In comparison to PRC2, the function of ASH1L in neuronal development is largely understudied. We report an essential role for ASH1L in neuronal morphogenesis. Using stem cell-derived human neurons, we show that acute knockdown of ASH1L in forebrain cortical excitatory neurons reduces neurite growth and arborization. One of the cellular mechanisms that underlie brain growth after birth is the development of neuronal arbors, and therefore failure to form connections due to reduced neuronal arborization could impair subsequent neuronal connectivity. The neuronal arborization defect in ASH1L deficient neurons correlates with growth cone-like structures that are enlarged. We propose that this regulation of neuronal growth is modulated by epigenetic counteracting mechanisms between ASH1L and PRC2, as we are able to rescue the ASH1L associated neuronal morphogenesis defects by inhibiting the activity of the PRC2 complex protein EZH2 (Laugesen et al., 2019; Schuettengruber et al., 2017). Furthermore, we uncover ASH1L as a novel regulator of the neurodevelopmentally important, BDNF-TrkB signaling pathway, which is essential for proper neuronal arborization during development (Gonzalez et al., 2016), as well as synapse development and function (Yoshii and Constantine-Paton, 2010). We show that the levels of NTRK2, the gene encoding the neurotrophin receptor TrkB, are significantly decreased in human neurons deficient for ASH1L compared to the other neurotrophin receptors. Downregulation of TrkB is sufficient to render the neurons unable to respond to exogenous BDNF, as the neuronal morphogenesis phenotypes are not rescued upon exposure to BDNF. Therefore, we propose that the deficit in TrkB expression caused by ASH1L knockdown decreases neuronal morphogenesis. Taken together, we present evidence of ASH1L’s modulation of neuronal morphogenesis by counteracting the activity of the PRC2 complex and by regulating the BDNF-TrkB signaling pathway.

## RESULTS

### ASH1L is dynamically expressed during human brain development

Because *ASH1L* is linked to ASD, using available databases we sought to determine the spatiotemporal expression pattern of ASH1L in the developing human brain, as a determinant of whether ASH1L could contribute to neuronal connectivity. Transcriptome data from human postmortem brain samples obtained from the Allen Brain Atlas (Hawrylycz et al., 2012), allowed us to assess the expression of *ASH1L* from fetal stages through adulthood (Fig. 1A and Supplementary Table S1 to S3). Temporally, *ASH1L* showed the highest expression between post conception weeks (pcw) 9 to 17, at year 1 postnatally, and from adolescence to adulthood (years 13 to 40 postnatally). Spatially, *ASH1L* showed the highest expression in prefrontal cortical (PFC) structures, including the dorsolateral, ventrolateral, and medial prefrontal cortex (DFC, VFC, MFC). Because ASH1L is an epigenetic regulator that generally activates transcription, we hypothesized that genes that are co-expressed at comparable levels in similar brain regions as *ASH1L* could provide some insight into the pathways modulated by ASH1L. Therefore, we used a correlation coefficient algorithm, which identifies the most highly correlated genes by highest coexpression with *ASH1L* within the three structures of the PFC, during the entire pre-natal window and then more specifically the DFC during 8 to 16 pcw (Supplementary Table S4). We then took the 500 most highly co-related genes for either the PFC or DFC and analyzed them with the *Toppgene* suite to determine if there are specific pathways or disease associations enriched within this gene cohort (Chen et al., 2009). These analyses showed enrichment in genes associated with intellectual disability and autistic disorders (Supplementary Fig. S1 and Supplementary Table S5-S6), along with an overrepresentation of pathways involved in neuronal morphogenesis, axon outgrowth, and synaptic function in both the DFC and the PFC. Similarly, these analyses showed an enrichment in genes associated with cytoskeletal regulation in the DFC and with chromatin regulation in the PFC (Fig. 1B, Supplementary Fig. S1 and Supplementary Table S5-S6). Together, these findings, along with the identification of ASH1L patients that have postnatal microcephaly, suggested that ASH1L could potentially have regulate neuronal arborization. We decided to further interrogate this possibility by modeling neuronal development using human stem cell derived cortical excitatory neurons (Fig. 1C) (Shi et al., 2012).

**Figure 1.**
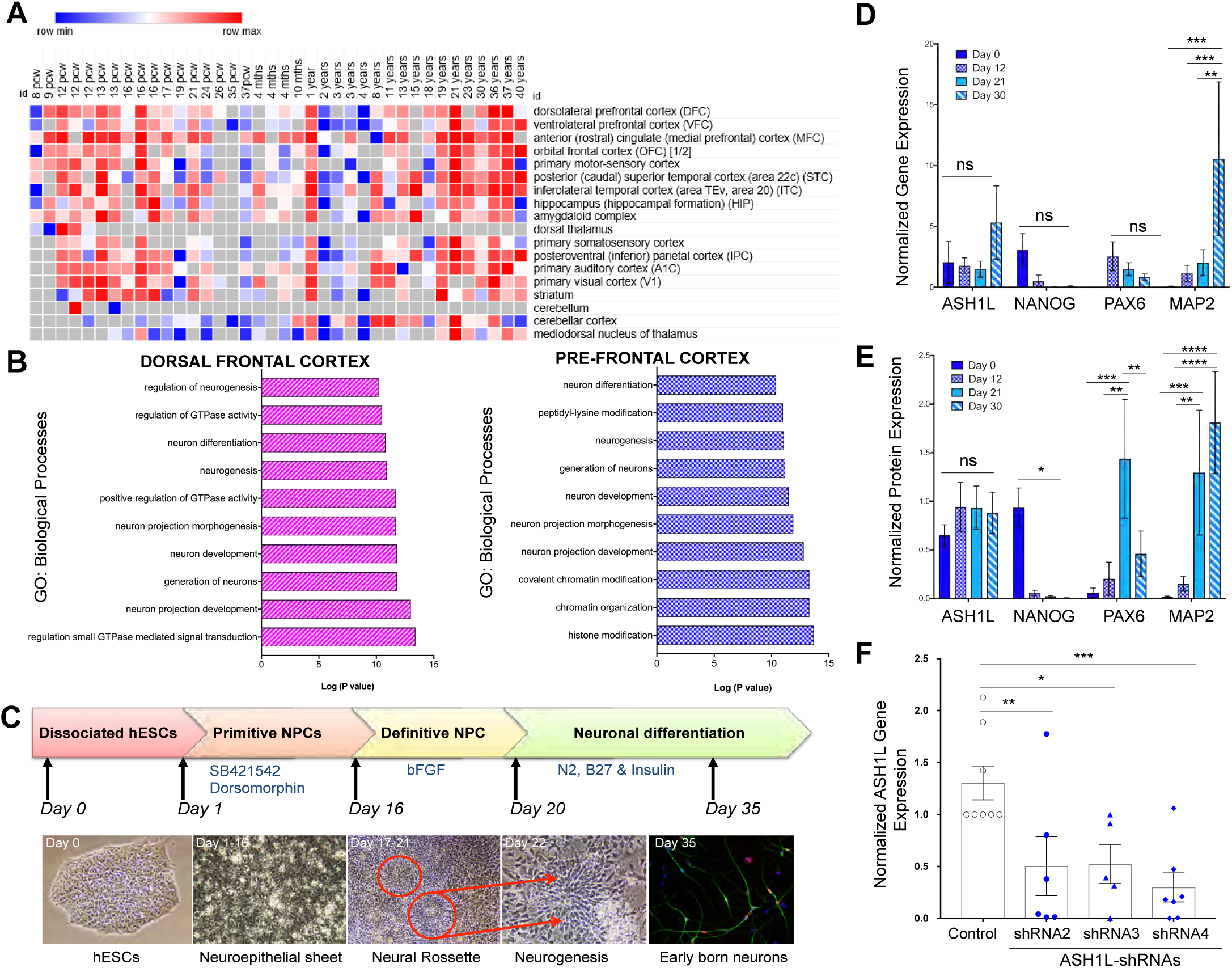
ASH1L is expressed throughout development *in vivo* and *in vitro*. (**A**) Analysis of ASH1L expression using the Allen Brain Atlas data set across 13 developmental stages and 8 to 16 brain structures. Heat map representation of ASH1L expression built using RNA-sequencing expression values downloaded from the developmental transcriptome database for ASH1L. Morpheus Software was used to build the Heat map showing each donor age with the corresponding brain region analyzed. Highest expression values are shown in red and lowest expression values are shown in blue. Grey values are used when no data was available for a particular developmental stage. (**B**) Pathway analysis of top-correlated genes with ASH1L during pre-natal development in DFC and PFC. Correlative genes with R values above 0.7 were identified between 8 to 37 pcw. GO categories for biological processes are shown as the Log10 of the adjusted p-value by Bonferroni correction. (**C**) Neuronal differentiation of hESCs into cortical deeper layer neurons. Top panel shows the major cellular stages from day 0 onwards during neuronal differentiation using a double SMAD inhibition protocol. Bottom panel shows corresponding images for each stage including hESCs colony before single cell dissociation (day 0), a neuroepithelial like sheet (days 1 to 16), the neural rosette stage (days 17-22), and deeper layer neurons (day 35). Deeper layer neurons were identified by co-immunostaining with CTIP2, a layer V marker (red) and β-III-Tubulin a pan-neuronal marker (green). (**D**) Analysis for ASH1L expression is shown at four distinct stages of in vitro neuronal differentiation for male H1 cells (days 0, 12, 21 and 30) in four independent experiments. Normalized gene expression is shown for *ASH1L*, *NANOG*, a stem cell marker, *PAX6*, a neuronal progenitor marker and *MAP2*, a mature neuron marker. Differential gene expression for MAP2 between day 0 to day 30 (***P = 0.0001), day 12 to day 30 (***P = 0.0001) and day 21 to day 30 (**P = 0.0013) by two-way ANOVA. (**E**) Protein expression analysis by western blot. Protein levels for ASH1L, NANOG, PAX6 and MAP2 were analyzed in 4 independent experiments. ASH1L protein levels were not significantly (ns) altered between different. NANOG protein expression was significantly different between day 0 and days 12, 21 and 30 (*P < 0.016). PAX6 protein expression varied between Day 0 to Day 21 (***P = 0.0003), Day 12 to Day 21 (**P = 0.0011) and Day 21 to Day 30 (**P = 0.0078). MAP2 protein was significantly differentially expressed between Day 0 and Day 21 (***P = 0.0007), Day 0 and Day 30 (****P < 0.0001), Day 12 to Day 21 (**P = 0.0022), and Day 12 to Day 21 (****P < 0.0001). Statistical analysis was carried using two-way ANOVA for multiple comparisons without corrections. (**F**) Gene expression analysis of *ASH1L* by qPCR in control or knockdown conditions. Expression of *ASH1L* normalized to *GAPDH* expression is shown for each experiment. The average of three technical replicates is shown per experiment. Control GFP (solid blue bar) = 1.31 ± 0.163, *n* = 8; shRNA-2 (solid light blue bar) = 0.504 ± 0.284, *n* = 6; shRNA-3 (hashed light blue bar) = 0.525 ± 0.189, *n* = 5; shRNA-4 (crisscrossed light blue bar) = 0.2989 ± 0.1402, n = 7. Ordinary one-way ANOVA uncorrected for multiple comparisons was used to obtain the p-values for: GFP vs. shRNA-2 (**P = 0.0069), GFP vs. shRNA3 (***P = 0.012) and GFP vs. shRNA4 (***P = 0.0008). In all graphs, results are shown as the mean ± standard error of the mean (SEM).

Autism has a higher incidence in males than females (Kim et al., 2011). To date the majority of *ASH1L* mutations have been reported in males with variable phenotypes (De Rubeis et al., 2014; Guo et al., 2018; Okamoto et al., 2017; Shen et al., 2019; Stessman et al., 2017; Wang et al., 2016). Therefore, for our studies we used stem cell derived human male neurons. We differentiated WA01 (H1) male human embryonic stem cells (hESCs) (Thomson et al., 1998), using a double SMAD inhibition protocol (Shi et al., 2012), to derive forebrain cortical excitatory neurons. We analyzed ASH1L gene (Fig. 1D) and protein (Fig. 1E and Supplementary Figure S2A) expression at day 0 for stem cells, at day 12 for neuroepithelial cells, at day 21 for neuronal progenitor cells, just as neurogenesis starts in these cultures, and at day 30 when deeper layer cortical excitatory neurons arise *in vitro*. We confirmed the in vitro developmental stage of our cultures using the following markers: NANOG for stem cells, PAX6 for neuronal progenitors, and MAP2 for mature neurons in four independent experiments (Fig. 1D-E). There were no significant differences in the *ASH1L* mRNA levels across the four *in vitro* developmental stages (day 0, 12, 21 & 30). *PAX6* and *NANOG* mRNA levels decreased as differentiation progressed and *MAP2* mRNA levels increased at the neuronal stage (day 30), showing appropriate neural induction and neuronal maturation of the cultures (Fig. 1D). Analysis of protein expression showed that ASH1L protein is expressed throughout these *in vitro* developmental stages, while NANOG was most highly expressed at stem cell stage (day 0), PAX6 was highly expressed at the neuronal progenitor stage (day 21), and MAP2 protein was present at highest levels at the neuronal stage (day 30) (Fig. 1E). Taken together these findings suggest that ASH1L function is important at multiple stages of development, which correlates with the embryonic lethality of *Ash1l ^-/-^* embryos (Zhu et al., 2016b).

### ASH1L regulates neuronal arbor and growth cone development

Patients with *ASH1L* mutations present with autistic behaviors, intellectual disability, and postnatal microcephaly, or failure of the brain to grow after birth (Okamoto et al., 2017). Postnatal microcephaly could arise from defects in neuronal axonogenesis, axonal and dendritic arborization, or synaptogenesis. While a role for ASH1L in synaptic function has been described in mice (Zhu et al., 2016b), mutant homozygous ASH1L animals are embryonic lethal and heterozygotes animals did not present any obvious structural brain phenotypes (Zhu et al., 2016b). Based on the pathways we identified in the correlation analysis (Fig. 1 and Supplementary Fig. S1) and the human patient phenotypes, we hypothesized that deficits in ASH1L in humans might compromise neurite outgrowth. To address this, we first tested five different shRNA constructs for knockdown of endogenous ASH1L in HEK293T cells and identified three shRNA constructs (shRNA2, 3 and 4) that consistently gave close to 80% knockdown of ASHL1 protein, after puromycin selection compared to control (Supplementary Fig. S2B-C and Supplementary Table S7). GFP control construct alone or ASH1L-shRNA plus a GFP construct were transfected into male cortical neurons (Fig. 1F and Supplementary Fig. S3). For this and all subsequent shRNA experiments, the shRNA transfected neurons were treated with puromycin for 72 hours, to enrich for transfected cells. shRNA transfection of these cortical neurons significantly reduced ASH1L gene expression levels by an average of 65.2% (Fig. 1E), compared to control transfected neurons, with no significant differences between the different shRNA transfected neurons.

Next, we determined whether acute knockdown of *ASH1L* could affect neuronal morphogenesis. For these experiments cortical neurons were co-transfected with a GFP construct plus either a control Luciferase-shRNA construct or the ASH1L-shRNA construct. After selection, neurons were immunostained with the pan-neuronal marker β-III-Tubulin, to confirm that GFP positive cells were neuronal-like cells (Fig. 2A). Morphometric analyses showed a 40.5% significant decrease in mean neurite length in male cortical neurons transfected with ASH1L-shRNA compared to controls. Similarly, we find a significant reduction of 59.5% in mean branching complexity index (measure of arborization) in ASH1L knockdown neurons compared to control neurons (Fig. 2B-C). Thus, in human male neurons ASH1L depletion decreases neurite outgrowth and arborization, suggesting that ASH1L modulates molecular mechanisms governing neuronal morphogenesis. Since we observed similar effects in neurite outgrowth with all ASH1L-shRNA constructs, and we find that for the branching complexity index is significantly smaller in ASH1L-shRNA2 neurons, all subsequent experiments were carried out using the ASH1L-shRNA2 construct (ASH1L-shRNA).

**Figure 2.**
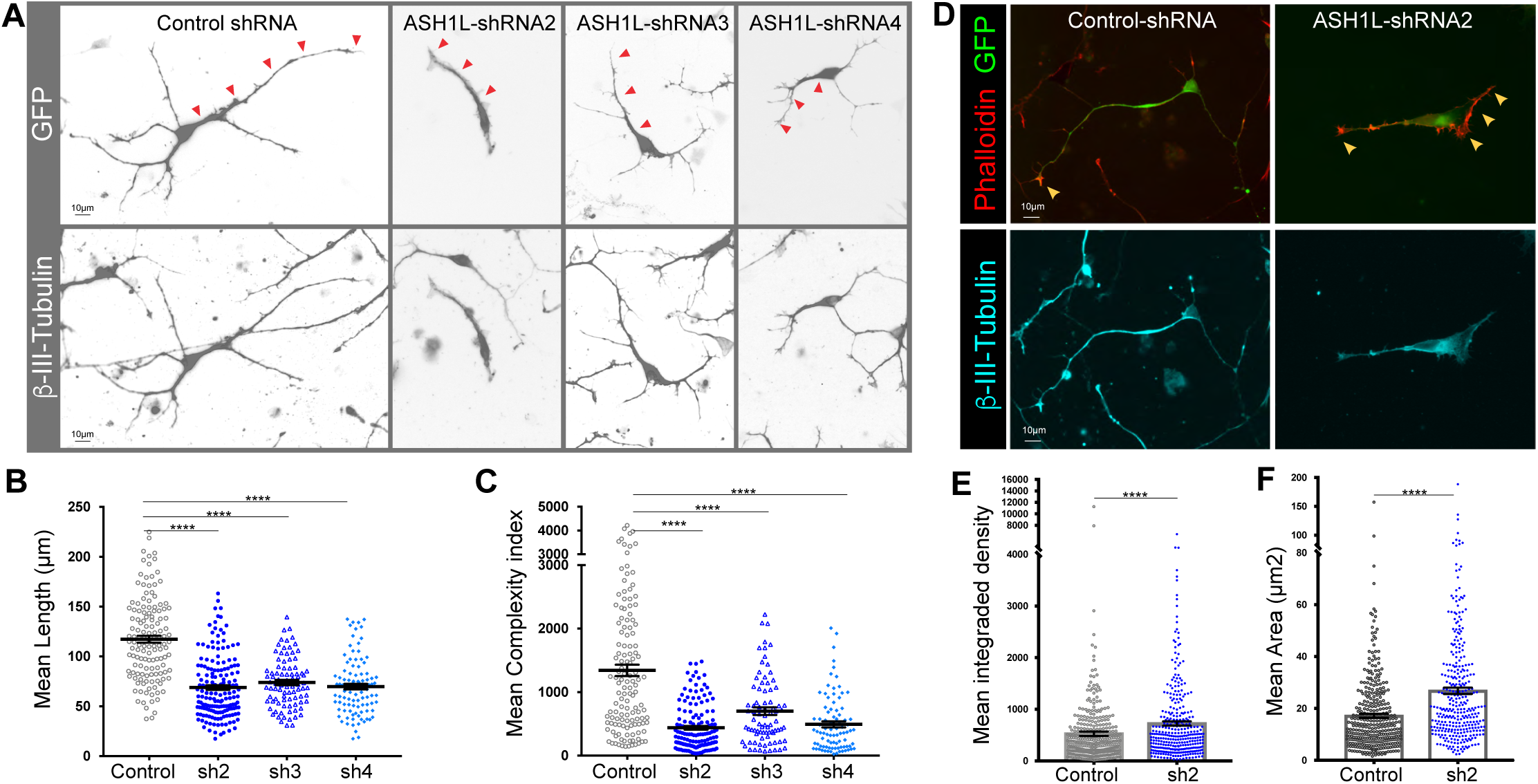
ASH1L modulates neurite outgrowth, branching, and growth cone size in cortical excitatory deeper layer neurons. (**A**) Defects in neurite length are observed upon knockdown of ASH1L in cortical excitatory neurons. Representative images are shown for neurons electroporated with GFP and control or ASH1L-targeting shRNA constructs. Top panels show GFP positive cells and bottom panels show β−III-Tubulin positive cells. For ease of viewing, grey scale images are shown as inverted images. Red arrows follow the longest neurite in the GFP positive neurons. Scale bar, 10µm. Neurites stained with β−III-Tubulin were measured using Neurolucida tracing software and no distinction was made between dendrites and axons. (**B**) Mean neurite length in cortical neurons under control or ASH1L knockdown conditions. Control shRNA (open grey circles) = 119.1 ± 3.92, *n* = 137 cells, N= 3 experiments; shRNA-2 (blue solid circles) = 74.87 ± 3.44, *n* = 177 cells, N= 5 experiments; shRNA-3 (open blue triangles) = 80.69 ± 4.28, *n* = 91 cells, N = 3 experiments; shRNA-4 (solid light blue diamonds) = 73.2 ± 3.3, *n* = 105 cells, N = 3 experiments. Statistical analysis was conducted using uncorrected Fisher’s test for multiple comparisons with ordinary one-way ANOVA (\*\*\*\**p* < *0.0001*). (**C**) Neurite branching complexity index in control and ASH1L knockdown. Control shRNA (open grey circles) = 1343 ± 89.46, *n* = 132 cells, N= 3 experiments; shRNA-2 (blue solid circles) = 438.5 ± 29.32, *n* = 150 cells, N= 5 experiments; shRNA-3 (open blue triangles) = 700.7 ± 58.45, *n* = 82 cells, N = 3 experiments; shRNA-4 (solid light blue diamonds) = 493.6 ± 47.07, *n* = 88 cells, N = 3 experiments. shRNA2 Vs. shRNA 3 ** P < 0.005; shRNA3 Vs. shRNA4 * P < 0.05. Statistical analysis was conducted using uncorrected multiple comparisons with ordinary one-way ANOVA (****P < 0.0001). All measurements were first analyzed for outliers using the ROUT (Q = 1%) method in all conditions and the analysis shown here was conducted after removal of outliers if present. In all graphs, error bars represent SEM, middle lane represents the mean value and all cell measurements are shown per condition for all experiments. (**D**) Enlarged growth cones are observed in neurons knockdown for ASH1L. Representative images are shown for neurons electroporated with GFP and control or ASH1L-targeting shRNA2 (sh2). Top panels show GFP positive cells (green) co-stained with phalloidin (red) and bottom panels show β−III-Tubulin positive cells (cyan). Scale bar, 10µm. Growth cone areas were measured using ImageJ**. (E)** Mean integrated density of phalloidin stained areas in control and ASH1L knockdown (shRNA-2) (N= 5 experiments). Control shRNA (open grey circles) = 526.8 ± 43.09, n=359 growth cones; shRNA-2 (blue solid circles) = 722.8 ± 42.31, n=331 growth cones. Statistical analysis was conducted using unpaired T-test with Welch’s correction (**P < 0.0015). **(F)** Mean Growth Cone Area (in µm*^2^*) in control and ASH1L knockdown (N=5 experiments). Control shRNA (open grey circles) = 17.18 ± 0.80, n=468 growth cones; shRNA-2 (blue solid circles) = 26.8 ± 1.21, n=337 growth cones. Statistical analysis was conducted using unpaired T-test with Welch’s correction (****P < 0.0001).

Our initial analysis of neuronal arbors also uncovered the presence of unusual growth cone-like structures in the ASH1L knockdown neurons. We identify the growth cones using phalloidin staining in combination with β-III-Tubulin staining in human neurons (Fig. 2D). Analysis of the integrated density in the growth cones showed a significant 28% increase in phalloidin staining under ASH1L knockdown conditions compared to controls (Fig. 2E). Comparably, analyses of the growth cone area showed a significant increase upon ASH1L knockdown (Fig. 2F). Reduction in neurite growth has been correlated with growth cone enlargement in *Aplysia* neurons for example (Ren and Suter, 2016), so the enlargement of the growth cone with ASH1L knockdown could reflect the reduced neurite length observed in human neurons.

### PRC2 and ASH1L counteracting activities modulate neuronal morphogenesis

We sought to determine the molecular mechanisms that could lead to neuronal morphogenesis defects associated with ASH1L knockdown by focusing on the well-known counteracting activities of Trithorax and Polycomb genes (Schuettengruber et al., 2017). ASH1L is a member of the Trithorax group of chromatin regulatory proteins. Trithorax proteins are chromatin modifying factors associated with gene activation during development (Miyazaki et al., 2013) and are proposed to modulate gene expression by counteracting Polycomb complex function (Supplementary Fig. S3-A) (Schuettengruber et al., 2017). PRC2 is a major chromatin regulator Polycomb complex that modifies chromatin by tri-methylation of histone H3 in Lysine 27 (H3K27me3), which is a chromatin repressive mark (Albert et al., 2017). In murine neurons, inhibition of EZH2, the catalytic subunit of the core PRC2 complex, increases dendritic arborization in hippocampal excitatory neurons (Qi et al., 2014). We hypothesized that PRC2 and ASH1L opposing activities might be regulating gene networks that modulate neuronal morphogenesis in human neurons. Thus, we asked whether impairing the catalytic activity of PRC2 in human neurons would counteract the effect of ASH1L knockdown (Supplementary Fig. S3-B). We exposed ASH1L-shRNA transfected human neurons to the EZH2 catalytic inhibitor - EI1 at 1.25 and 2.5 µM for 3 or 5 days in culture (Fig. 3A-C and Supplementary Fig. S3-C to D). After 3 days of EI1 treatment, there was a significant but modest rescue in neurite length with 2.5 µM EI1 (Fig. 3B), but after 5 days a there was a significant increase in neurite length with both 1.25 and 2.5 µM EI1, to levels comparable to the untreated control cells (Fig. 3C). Analysis of neurite branching showed no significant rescue after 3 days with either dosage of EI1 (Supplementary Fig. S3C). However, ASH1L knockdown neurons treated with 1.25 µM EI1 for 5 days showed a modest but significant rescue in the number of branches per cell that was significantly different from the untreated ASH1L-shRNA neurons (Supplementary Fig. S3D). Therefore, inhibition of the catalytic activity of PRC2 primarily rescues the neurite length phenotype in ASH1L-knockdown neurons and more modestly the arborization phenotype. Taken together these findings suggest that there is antagonism between the Trithorax protein ASH1L and the PRC2 complex proteins during the development of neuronal connectivity with neuronal morphogenesis, being determined by this balance.

**Figure 3.**
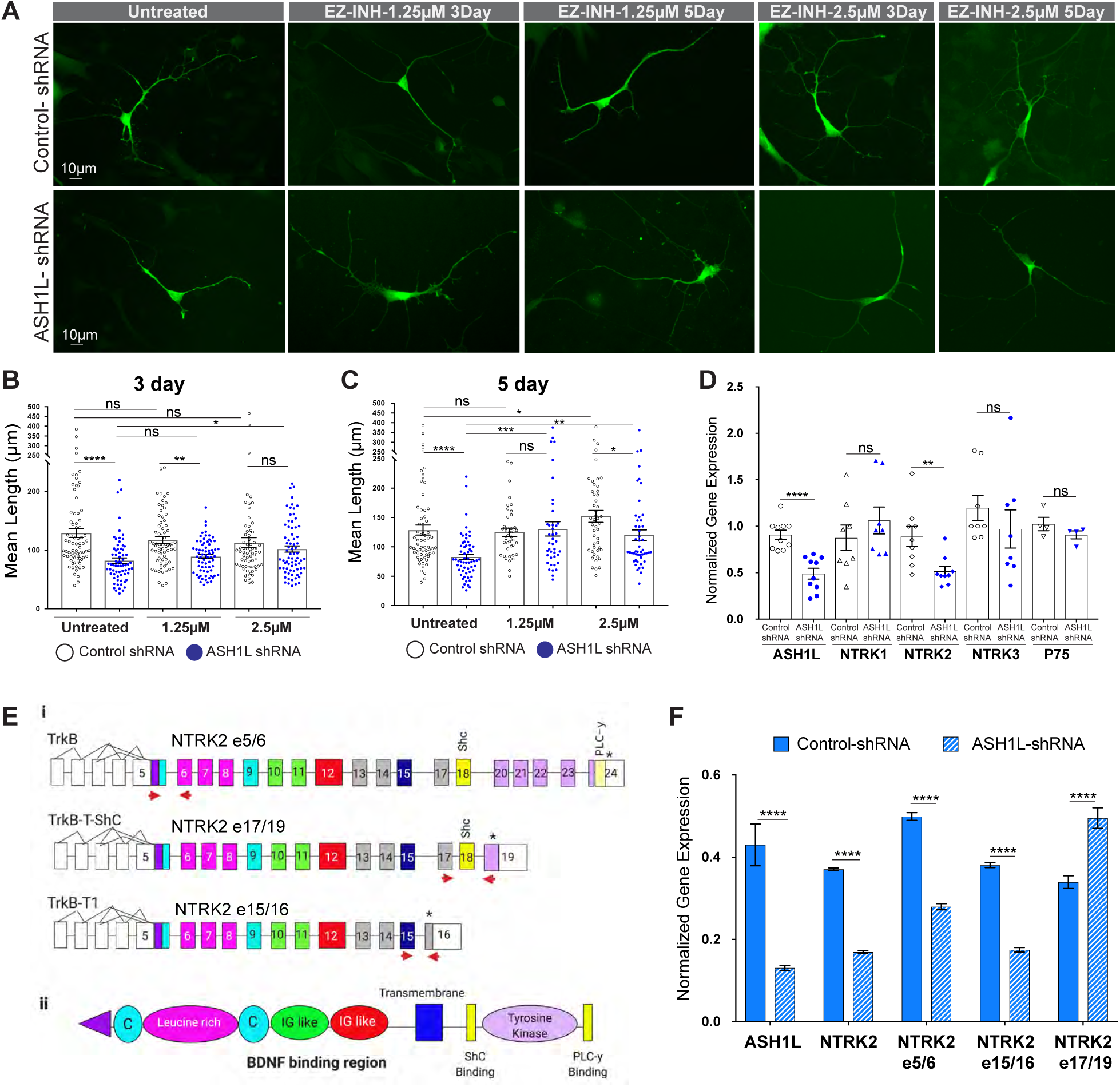
ASH1L phenotype is rescued by EZH2 inhibition and correlates with downregulation of NTRK2. (**A**) Representative images are shown for neurons electroporated with GFP and either control-shRNA or ASH1L-shRNA constructs treated with 1.25µM or 2.5µM EZH2 inhibitor (EI1) for 3 and 5 days. Scale bars, 10µm. (**B**) Mean neurite length in cortical neurons under control or ASH1L knockdown conditions were either untreated or treated for 3 days with 1.25µM or 2.5µM EI1. All measurements per cell are shown for each condition in combination with the mean ± SEM. All control shRNA are open black circles and ASH1L-shRNA are solid blue circles. Control shRNA = 129.2 ± 7.78, n = 74 cells *vs* ASH1L-shRNA = 82.2 ± 4.62, n = 69 cells; Control-shRNA+1.25µM EI1 = 117.1 ± 5.43, n = 69 cells *vs* ASH1L-shRNA +1.25µM EI1 = 88.9 ± 3.41, n = 70 cells; Control-shRNA + 2.5µM EI1 = 112.6 ± 8.64, n = 67 cells *vs* ASH1L-shRNA + 2.5µM EI1 = 101.9 ± 5.30, n = 73 cells. Statistical analysis was conducted using multiple comparisons with ordinary one-way ANOVA uncorrected for multiple comparisons. ****P < 0.0001; **P < 0.0015, *P < 0.025, and not significant (ns). (**C**) Mean neurite length in cortical neurons under control or ASH1L knockdown conditions were either untreated or treated for 5 days with 1.25µM or 2.5µM EI1. All control shRNA are open black circles and ASH1L-shRNA are solid blue circles. All measurements per cell are shown for each condition in combination with the mean ± SEM. Control-shRNA= 128.5 ± 8.53, n = 64 cells; ASH1L-shRNA = 82.9 ± 4.87, n = 59 cells; Control-shRNA+1.25µM EI1 = 124.7 ± 6.99, n = 40 cells; ASH1L-shRNA+1.25µM EI1 = 130.7 ± 12.22, n = 41 cells; Control-shRNA + 2.5µM EI1 = 151.9 ± 10.16, n = 48 cells; ASH1L-shRNA + 2.5µM EI1 = 120 ± 8.89, n = 50 cells. Statistical analysis was conducted using multiple comparisons with ordinary one-way ANOVA uncorrected for multiple comparisons. ****P < 0.0001, ***P < 0.0003, **P < 0.002, *P < 0.05, and not significant (ns). (**D**) Analysis of gene expression across genes encoding the different neurotrophin receptors in control and ASH1L knockdown cortical neurons. qPCR analysis of expression shown as normalized data to reference gene GAPDH that was then normalized to control is shown as normalized ΔΔCt values for control (black open icons) and ASH1L shRNA (blue solid icons) samples for the following genes: ASH1L (n=16 independent experiments, ****P <0.0001); NTRK1 (n = 11 independent experiments, ns); NTRK2 (n=14 independent experiments, **P < 0.0013); NTRK3 (n = 11 independent experiments, ns); and P75 (n=5 independent experiments, ns). Statistical analysis was carried out using paired t-tests for each gene. (**E**) Schematic representation of TrkB isoforms analyzed. Top panel (i) shows the splicing isoforms of TrkB gene containing the N-terminus (NTRK2 e5/6). Full length TrkB and isoforms TrkB-T-Shc (NTRK2 e17/19) and TrkB-T1 (NTRK2 e15/16). Red arrows show the location of the primers utilized to amplify the specific isoforms. Bottom panel (ii) shows the full length TrkB proteins showing the corresponding protein domains to corresponding exons by color coding. Adapted from (Luberg et al., 2010). (**F**) Analysis of NTRK2 isoform expression by ddPCR in male cortical neurons under control or ASH1L knockdown conditions. Data was normalized to GAPDH and then normalized to control and is shown as counts/ng of DNA for three independent experiments. Solid blue bars are control shRNA samples while hashed blue and white bars are ASH1L-shRNA samples for the following transcripts: *ASH1L* (Control-shRNA = 0.430 ± 0.088 *vs.* ASH1L-shRNA = 0.131 ± 0.010; ****P< 0.0001); *NTRK2* full length (control-shRNA = 0.371 ± 0.005 *vs.* ASH1L-shRNA = 0.170 ± 0.006; ****P < 0.0001); *NTRK2 e5/6* (control-shRNA = 0.499 ± 0.05 *vs.* ASH1L-shRNA=0.279 ± 0.013; ****P< 0.0001); *NTRK2 e15/16* (control-shRNA = 0.381 ± 0.010 *vs.* ASH1L-shRNA=0.175 ± 0.010; ****P< 0.0001); and *NTRK2 e17/19* (control-shRNA = 0.339 ± 0.027 *vs.* ASH1L-shRNA=0.495 ± 0.043; ****P < 0.0001).

### ASH1L specifically regulates the expression of the neurotrophin receptor TrkB and its isoforms

The antagonizing activities of ASH1L and PRC2 could regulate gene programs important for neuronal morphogenesis. Neurotrophin-signaling is essential for neuronal arborization and neurite outgrowth (Deinhardt and Chao, 2014a). We used the ENCODE database to determine if epigenetic marks associated with transcriptional activation or repression are seen at human genes for the neurotrophin receptors, *NTRK1* (TrkA), *NTRK2* (TrkB), *NTRK3* (TrkC) and *P75* (P75) (Deinhardt and Chao, 2014b). Using data available for female H9 stem cell derived cortical excitatory neurons (as this was the only dataset available for cortical neurons), we aligned each gene to epigenetic marks associated with transcriptional repression by PRC2 (H3K27me3) (van Mierlo et al., 2019), and associated with active gene transcription (H3K36me3 and H3K4me3) (Balbach and Orkin, 2016; Eram et al., 2015; Shao et al., 2014). The current ENCODE data sets lack analysis of H3K36me2 that is primarily deposited by ASH1L. However, H3K36me2 presence will lead to upregulation of H3K36me3 by SET2 (Li et al., 2019a), and H3K4me3 marks are deposited by the ASH1L interactor - MLL complex (Zhu et al., 2016a). Therefore, both H3K36me3 and H3K4me3 marks are highly correlated with the presence of H3K36me2 (Miao and Natarajan, 2005), and were used as correlative evidence for the presence of H3K36me2 marks. We found that NTRK1 is almost entirely enriched for the PRC2 associated H3K27me3 epigenetic marks along the gene body. Similarly, but to a lesser extent, NTRK3 is enriched in H3K27me3 histone marks along the gene body, while P75 shows high enrichment in H3K27me3 marks at its transcription start site. In contrast, NTRK2 is primarily enriched in H3K36me3 along the gene body and H3K4me3 closer to the promoter regions (Supplementary Fig. S3-E). These results suggest that at least in H9 cortical excitatory neurons NTRK1, NTRK3 and P75 expression could be suppressed by PRC2, while NTRK2 is being actively transcribed in cortical neurons. To determine if ASH1L could differentially modulate the expression of the different neurotrophin receptors, we evaluated neurotrophin receptor mRNA levels using reverse-transcription-coupled quantitative PCR (RT-qPCR) analyses on male cortical neurons that were transfected with control and ASH1L-shRNA constructs. Knockdown of ASH1L in cortical neurons is associated with a 41.8% significant decrease in NTRK2 mRNA, but NTRK1, NTRK3 and P75 mRNA levels were not significantly affected (Fig. 3D). Since, binding of BDNF to its TrkB receptor is essential for neuronal arborization (Segal et al., 1995), this raises the possibility that the epigenetic regulation of TrkB by ASH1L could modulate neuronal morphogenesis. The PCR primers used for NTRK2 were designed to detect all known splice variants of this gene. Since TrkB has several functional variants generated by alternative splicing, we next evaluated expression of the NTRK2 splice variants in the ASH1L knockdown neurons.

Histone H3K36 methylation regulates pre-mRNA splicing in *S. cerevisiae* (Sorenson et al., 2016), and MRG15, which forms a complex with ASH1L (Lee et al., 2019), also regulates pre-mRNA splicing (Iwamori et al., 2016). Since ASH1L dimethylates H3K36 (Miyazaki et al., 2013; Zhu et al., 2016a) and forms a complex with MRG15, we asked if ASH1L could regulate the alternative splicing of NTRK2 (TrkB) mRNA, which has seven distinct splicing isoforms. Four of these isoforms are expressed in the human PFC during postnatal development (Luberg et al., 2010). Because our cultures are composed of forebrain cortical neurons, we decided to focus on 3 isoforms expressed in the PFC for which their sequence was amenable to PCR based detection (Fig. 3E-F). Using digital droplet RT-PCR (dd-PCR) in control and ASH1L shRNA-transfected neurons, we found a significant decrease in expression of isoforms containing exons 5 and 6 (NTRK2 e5/6). These isoforms include the full length TrkB (NTRK2-FL) and all isoforms that contain the N-terminus of the TrkB protein. Two additional isoforms that we examined were TrkB-T1 (NTRK2 e15/16) (Fryer et al., 1996) and TrkB-SHC (NTRK2 e17/19) (Wong and Garner, 2012). TrkB-T1 lacks the tyrosine kinase domain, SHC, and PLCγ binding domains. TrkB-T1 acts as a dominant negative that scavenges the full length TrkB, leading to decreased BDNF-TrkB signaling (Luberg et al., 2010). TrkB-SHC contains the SHC binding domain, but lacks the PLCγ binding domain and the tyrosine kinase domain, and has been proposed to decrease the levels of phosphorylated TrkB (Luberg et al., 2010; Wong and Garner, 2012). We found a significant downregulation of isoform NTRK2 e15/16, which encodes TrkB-T1. However, we found significant upregulation of NTRK2 e17/19, which encodes TrkB-SHC. Mice with a loss of function TrkB-T1 have reduced dendritic complexity (Carim-Todd et al., 2009). Taken together these data suggest that ASH1L modulates the expression of full length TrkB and potentially two other N-terminus containing isoforms, TrkB-T1 and TrkB-SHC, which could further exacerbate the neuronal morphogenesis phenotype.

### Exogenous addition of BDNF neurotrophin does not rescue ASH1L neuronal phenotypes and highlights an intact albeit dampened downstream signaling cascade

BDNF binding to TrkB receptor will elicit the endocytosis of the receptor bound ligand and in turn elicit a signaling cascade that leads to the phosphorylation and subsequent nuclear translocation of the CREB transcription factor. CREB activation promotes the transcription of both BDNF and TrkB (Deogracias et al., 2004; Finkbeiner, 2000a, b). Additionally, BDNF has been shown to elicit local translation in dendritic and axonal compartments (Leal et al., 2014). Human stem cell-derived neurons are known to express BDNF in culture, but with decreased TrkB expression, the ASH1L depleted neurons may not be able to respond to the endogenously expressed BDNF. Thus, we asked if the endogenous BDNF levels are sufficient to activate the lower TrkB levels in the ASH1L depleted human cortical neurons by exogenously adding BDNF to our cultures. We tested the possibility that exogenous BDNF at receptor saturating levels (10 ng/ml) might rescue the neurite growth deficit seen with ASH1L depletion (Supplementary Fig. S4-A). In control neurons, we find that after 96 hours of exposure to exogenously added BDNF, there is a significant 15% increase in mean neurite length, and a 35% increase in mean branch number, indicating that the TrkB receptors are not saturated with endogenous BDNF (Fig. 4A-C). However, in ASH1L knockdown neurons the addition of exogenous BDNF caused a very modest increase in mean neurite length and branching that was not statistically different from ASH1L knockdown neurons that were not treated with BDNF (Fig. 4A-C). This finding posited the question of whether the downstream signaling pathway remains intact after downregulation of ASH1L.

**Figure 4.**
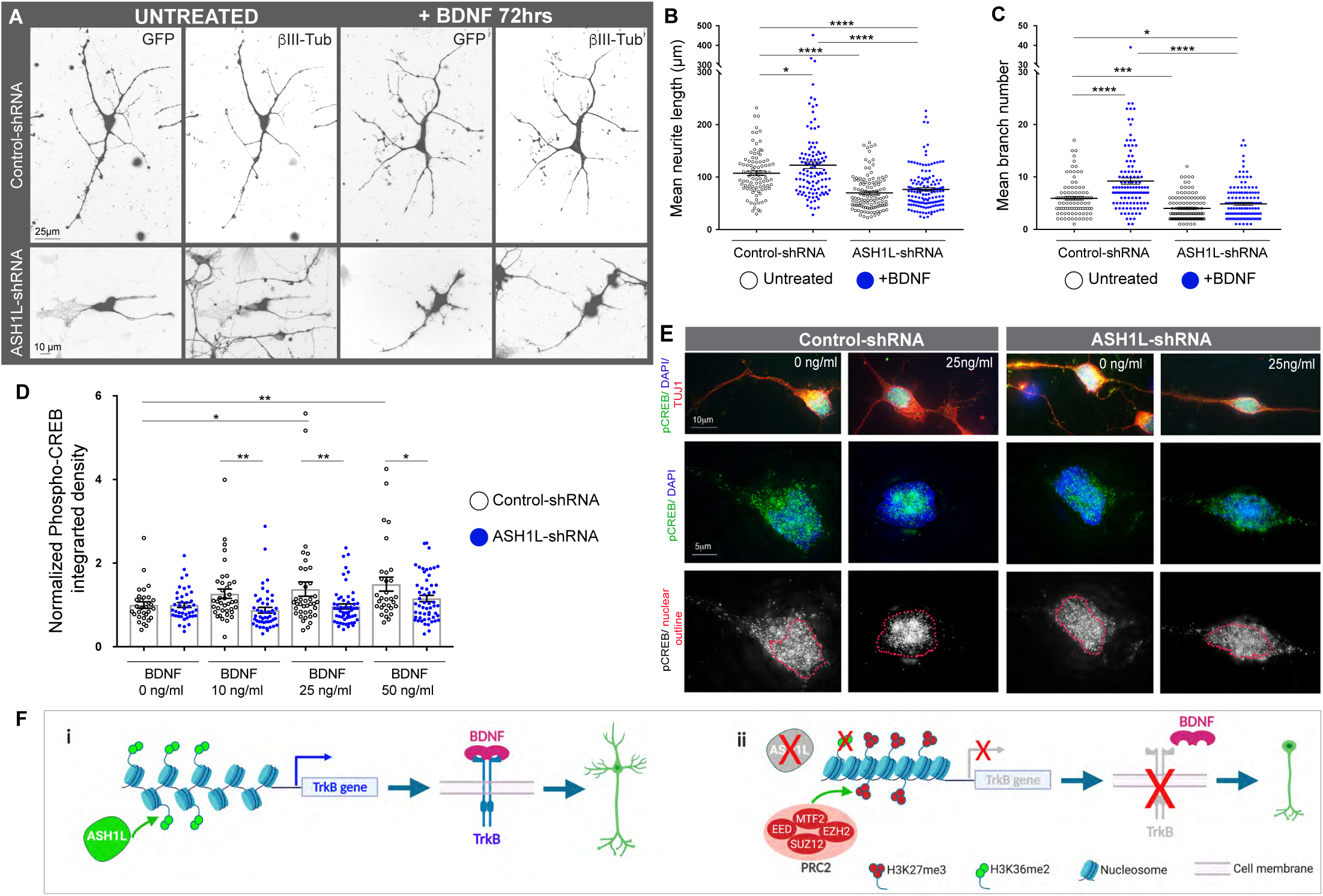
Loss of ASH1L reduces CREB activation by BDNF. (**A**) Representative images are shown for neurons electroporated with GFP and either control shRNA or ASH1L shRNA constructs. Neurons treated with 0 or 10 ng/ml of BDNF for 3 days after puromycin selection positive for GFP and with □-III-Tubulin (□-III-Tub) shown. Scale bar, 25µm for controls and 10µm for ASH1L-shRNA samples. For ease of viewing images were processed from RGB to luminescence in ImageJ and then inverted. (**B**) Mean neurite length in cortical neurons under control or ASH1L knockdown with or without 72hr BDNF treatment. Untreated samples are shown in open black circles and BDNF treated samples are shown in solid blue circles. Control shRNA = 107.3 ± 4.36, n = 93 cells; Control shRNA +BDNF = 122.7 ± 6.29, n = 111 cells; ASH1L-shRNA = 69.89 ± 3.179, n = 112 cells; and ASH1L-shRNA +BDNF= 76.46 ± 3.15, *n* = 144 cells. ****P < 0.0001, and * P < 0.02. If no p-values are shown that denotes that the differences were not significant. (**C**) Mean branch number is shown for cortical neurons under control or ASH1L-shRNA knockdown with or without 72hr BDNF treatment. Untreated samples are shown in open black circles and BDNF treated samples are shown in solid blue circles. Control shRNA = 5.93 ± 0.34; Control shRNA +BDNF = 9.22 ± 0.57; ASH1L-shRNA = 4.01 ± 0.23; and ASH1L-shRNA +BDNF = 4.86 ± 0.27. ***P < 0.0001, ***P < 0.0006, and * P <0.045. If no p-values are shown that denotes that the differences were not significant. All measurements are shown per cell for each condition for four independent experiments. Mean is shown as the middle line and error bars are SEM. Statistical analysis was conducted using uncorrected Fisher’s test for multiple comparisons with ordinary one-way ANOVA. **(D)** Activation of phospho–CREB is reduced in ASH1L knockdown neurons. Control or ASH1L shRNA transfected neurons were acutely exposed for 15 minutes to increasing doses of BDNF (0, 10, 25 and 50 ng/ml) to measure the nuclear translocation of phospho-CREB. Analysis of phospho-CREB nuclear levels was conducted by measuring integrated density on nuclear region of interest (ROI) after background subtraction in images acquired in a widefield fluorescent microscope. All values were normalized to the mean average value of the BDNF 0ng/ml samples for each experiment both in control-shRNA (open black circle) and ASH1L-shRNA (solid blue circle). Control-shRNA + 0ng/ml BDNF =1 ± 0.077, n=31 cells; *Vs*. ASH1L-shRNA + 0ng/ml BDNF = 1 ± 0.060, n=41 cells. Control-shRNA +10ng/ml BDNF = 1.27 ± 0.111, n=38 cells; *Vs*. ASH1L-shRNA + 10ng/ml BDNF = 0.87 ± 0.069, n=48 cells, ** P < 0.006. Control-shRNA +25ng/ml BDNF = 1.379 ± 0.171, n=39 cells; *Vs*. ASH1L-shRNA + 25ng/ml BDNF = 0.9665 ± 0.060, n=55 cells, ** P < 0.0025. Control-shRNA +50ng/ml BDNF = 1.498 ± 0.166, n=31; *Vs*. ASH1L-shRNA + 50ng/ml BDNF = 1.15 ± 0.074, n=55 cells, * P < 0.02. Control-shRNA + 0ng/ml BDNF *Vs*. Control-shRNA +10ng/ml BDNF. Control-shRNA + 0ng/ml BDNF *Vs*. Control-shRNA +25ng/ml BDNF, * P < 0.02. Control-shRNA + 0ng/ml BDNF *Vs*. Control-shRNA +50ng/ml BDNF, ** P < 0.003. If no p-values are shown that denotes that the differences were not significant. All values are shown as mean ± SEM. All statistical analysis was carried out using uncorrected Fisher’s test for multiple comparisons in one-way ANOVA on all data points for three independent experiments. (**E**) Super resolution microscopy confirms increased nuclear phospho-CREB signal in control neurons compared to ASH1L knockdown neurons in response to exogenous BDNF. Representative images are shown for neurons transfected with either control-shRNA or ASH1L-shRNA that were either untreated (0 ng/ml) or treated with BDNF (25ng/ml). Top panels show phospho-CREB (green) and TUJ1 (□-III-Tubulin) a pan-neuronal marker (red), in transfected neurons. Middle panels show co-immunostaining of phospho-CREB (green) and nuclear stain DAPI (blue). Bottom panels show the phospho-CREB signal in grey and the nucleus is outlined in red. (**F**) Model depicting epigenetic regulation by ASH1L and PRC2 of the BDNF-TrkB signaling pathway and influence in neuronal morphogenesis. In the top panel (i) ASH1L contributes to the expression of NTRK2 the gene encoding TrkB by demethylating H3K36 (green) and preventing repression by PRC2, leading to normal production of the TrkB receptor and normal neurogenesis. In the bottom panel (ii) reduction in ASH1L levels lead to reduce levels of H3K36me2 in the NTRK2 gene, allowing PRC2 to trimethylate H3K27 (red), leading to the repression of NTRK2, which results in the reduction of the levels of TrkB receptor.

BDNF-dependent activation of TrkB is known to induce activation and nuclear translation of the transcription factor CREB through its phosphorylation (Watson et al., 1999). Phosphorylated CREB is needed to activate transcription of genes for neurite outgrowth and neuronal morphogenesis. Thus, we reasoned that if TrkB signaling were intact in the ASH1L depleted neurons, BDNF stimulation would increase nuclear localization of phospho-CREB. shRNA transfected neurons were acutely treated with BDNF at increasing doses (0, 10, 25 and 50 ng/ml) after puromycin selection (Fig. 4C and Supplementary Fig. S4-B). Analysis of widefield fluorescence images showed a significant increase in nuclear phospho-CREB signal in control neurons treated with increased BDNF doses. However, in ASH1L knockdown neurons increased BDNF levels did not increase nuclear phospho-CREB signal compared to the untreated control (Fig.4D). We further confirmed these findings by using super resolution microscopy which showed a distinct nuclear localization for phospho-CREB within 15 minutes after 25 ng/ml BDNF exposure in control neurons compared to reduced nuclear localization of phospho-CREB in ASH1L knockdown neurons (Fig. 4E). We wondered if the decrease in phospho-CREB signal might be indicative of reduced expression of CREB itself. However, after knockdown of ASH1L in cortical neurons we found no significant change in mRNA levels of *CREB1,* the gene encoding CREB (Supplementary Fig.S4-C). Taken together, these findings show that CREB gene expression is maintained despite the knockdown of ASH1L, but the reduction of ASH1L results in decreased activation of CREB by BDNF.

## DISCUSSION

In mice, ASH1L is essential for embryonic development, as ASH1L^-/-^ mice are early embryonic lethal (Zhu et al., 2016b). While ASH1L^-/+^ mice had no major brain abnormalities reported (Zhu et al., 2016b), they did show changes in gene expression in response to neuronal activity (Zhu et al., 2016b). Furthermore, RNA interference of the *ASH1L* homologue (*Ash1)* in *Drosophila* disrupted their capacity to complete habituation tasks, suggesting a major disruption of neuronal function (Stessman et al., 2017). However, the extent to which ASH1L contributes to the development of human neuronal circuitry is largely unknown. We find that acute knockdown of ASH1L stunts neurite growth and reduces neurite branching in male human stem cell-derived cortical neurons. The reduced neurite outgrowth is accompanied by increased growth cone size, suggesting that reduction in ASH1L may cause reduced growth rates and could potentially lead to neuronal guidance defects *in vivo*. Our correlation analysis of the top co-expressing genes with ASH1L shows an overrepresentation of genes involved in neurite outgrowth and cytoskeletal function. In fact, we further identified that reducing the BDNF-TrkB signaling pathway by downregulation of gene encoding the TrkB receptor, *NTRK2*, is one potential mechanism underlying the ASH1L associated knockdown neuronal phenotypes. Furthermore, our acute BDNF signaling experiments showed that reduction in TrkB gene expression, leads to a reduced response to BDNF as downregulation of ASH1L leads to diminished levels of nuclear phospho-CREB signal. Since, the BDNF-TrkB signaling pathway is critical for neuronal arborization, our findings suggest that ASH1L is important for the early steps of neuronal circuitry development, including neuronal arborization. It is clear that, ASH1L has multiple targets as an epigenetic regulator, but our data indicate that its regulation of the BDNF-TrkB signaling pathway likely contributes to its role in neuronal morphogenesis.

*In vivo* and *in vitro* evidence suggest that ASH1L contributes to the methylation of H3K4 and H3K36 (Gregory et al., 2007; Xia et al., 2013). The dimethylation of H3K36 (H3K36me2) and trimethylation of H3K4 (H3K4me3) sites modify chromatin structure, allowing for transcriptional activation of target genes (Miyazaki et al., 2013; Zhu et al., 2016a). Methylation of H3K36 has been proposed to counteract the activity of the Polycomb group and prevent deposition of the transcriptionally repressive H3K27me3 mark (Schuettengruber et al., 2017). ASH1L deposition of the H3K36me2 mark has been reported to inhibit the catalytic activity of PRC2 (Huang and Zhu, 2018). This mechanism of opposing activities of the Polycomb and Trithorax complex is essential during organogenesis (Schuettengruber et al., 2017). For example, the PRC2 complex through the tri-methylation of H3K27 regulates cell fate transitions during corticogenesis (Pereira et al., 2010). In particular, deletion of EZH2, the PRC2 catalytic subunit (Schuettengruber et al., 2017) early in development leads to precocious neuronal differentiation and astrogliogenesis (Corley and Kroll, 2015). In addition to its role in cell fate determination, EZH2 has also been implicated in neuronal arborization, as knockdown of EZH2 results in increased branching of neuronal arbors in hippocampal neurons and modulated BDNF expression (Qi et al., 2014). We find that human PRC2 contributes to development of neuronal arbors, as inhibition of EZH2 rescues primarily the neurite outgrowth phenotypes associated with ASH1L knockdown. Based on this data, we hypothesize that in the absence of ASH1L, the activating H3K36me2 marks on NTRK2 are reduced, enabling PRC2 to place repressive H3K27me3 marks and reduce NTRK2 expression (Fig. 4F). Taken together, our findings point to the counteracting activities of ASH1L and PRC2 as modulators of gene networks associated with the development of neuronal connectivity.

Recently, ASH2L, a member of the Trithorax core complex “COMPASS”, was implicated in cell fate transitions during murine cortical development (Li et al., 2019b). Since we did not knockdown ASH1L at the neuronal progenitor stage our findings do not discount the potential of ASH1L to be involved in cell fate transitions during human corticogenesis. Additionally, we do not discount that additional epigenetic mechanisms could be at play such as the activity of specific histone demethylases that regulate the methylation of lysine 36 (Li et al., 2019a) and could facilitate the activity of the Polycomb complexes. Finally, while the current work has focused on the ASH1L-PRC2 axis, we are aware that ASH1L could also oppose the activity of the PRC1 complex during embryonic development. PRC1 also acts antagonistically to Trithorax protein complexes, and in *Drosophila,* the PRC1 complex regulates neuronal arborization of class IV sensory neurons (Parrish et al., 2007). Our results combined with these recent studies provide the initial mechanistic insights for understanding the role of epigenetic regulation of neuronal morphogenesis.

## MATERIALS & METHODS

### Allen Brain atlas analysis of ASH1L expression

Gene expression data was obtained from the Allen Brain Atlas Developmental Transcriptome Database (© 2010 Allen Institute for Brain Science. Allen Human Brain Atlas. Available from: <u>human.brain-map.org</u>). Transcriptome data was obtained from RNA-sequencing and exon microarray hybridization that was generated across 13 developmental stages and 8-16 brain structures. This data was previously obtained by RNA-sequencing that was performed in human brain specimens collected from normally developing individuals ranging from 8 pcw to over 40 years of age by the Department of Neurobiology at Yale School of Medicine and the National Institute of Mental Health (Miller et al., 2014). In total, expression data was analyzed from 37 different donors; 21 males and 16 females. Raw data is shown in supplementary materials (Supplementary Table S1-B).

Expression heat maps were generated using RNA-sequencing expression values downloaded from the Developmental Transcriptome database for genes of interest. Expression values, age of donor, and brain structure were further analyzed using the Broad Institute’s Morpheus software (https://software.broadinstitute.org/morpheus). In cases where multiple donors were identified for the same stage, the expression was averaged (12pcw, 13pcw, 16pcw, 4 months, 3 years) in order to improve the visualization of age-related expression trends (Supplementary Table S1-S2). Donor information can be found in (Supplementary Table S3). Average expression, hierarchical clustering, and box & whisker plots were calculated and created using the Morpheus software.

### Identification of ASH1L co-expressing genes and functional enrichment analysis

To identify genes with similar expression patterns to *ASH1L* and other genes of interest in the study, the correlative search option within the Developmental Transcriptome Database was used over a time frame that corresponds to prenatal development (8 to 37 pcw). We focused the correlative search on structures of the PFC and DFC and identified the top 500 genes with the highest positive correlation (r) to *ASH1L* gene expression (0.741<r < 0.959). To analyze the function of top correlated genes Toppgene software (https://toppgene.cchmc.org/enrichment.jsp) was used to detect functional enrichment of these top genes based on Transcriptome, Proteome, Ontologies (GO, Pathway), Phenotype (human disease and mouse phenotype), and literature co-citation (Chen et al., 2009). Genes were imported into the portal, correction was set and resulting functional enrichments were ranked by p-value.

### Stem cell culture

Human embryonic H1 (WiCell catalog # WAe0001-A, NIH approval: NIHhESC-10-0043) male stem cells (hESCs) and female H9 (WiCell catalog # WAe009-A, NIH approval: NIHhESC-10-0062) were used to analyze the role of ASH1L in neuronal development. hESCs were cultured as previously described (Ludwig et al., 2006). Briefly, cells were grown on matrigel (Corning catalog # 356234), a feeder-free substrate, and fed daily with complete changes of mTSER media (StemCell technologies catalog #85850). hESCs colonies were inspected daily and spontaneous differentiating colonies were manually removed. Cells were passaged every 4 to 7 days with ReLeSR reagent (StemCell Technologies catalog #05872) to ensure that only undifferentiated stem cells remained in the cultures. Stem cell cultures were passaged at a 1:6 ratio each time. For neuronal differentiation experiments, cells under 45 passages were used to minimize the chances of spontaneous chromosomal abnormalities that arise after passage number 50 (Thomson et al., 1998).

### Generation of forebrain cortical neurons from stem cells

hESCs were differentiated using a previously described double SMAD inhibition protocol that promotes the formation of the neuroectoderm by preventing the formation of mesoderm and endoderm (Shi et al., 2012). This protocol produces primarily excitatory forebrain cortical neurons in a monolayer and was conducted with slight modifications. In brief, hESCs were dissociated into single cells using Gentle Cell Dissociation Reagent (StemCell Technologies Catalog #07174). Dissociated hESCs were counted and 1.5 to 2 million cells were plated per well of a 12-well plate coated with matrigel (Corning Catalog #354277). Cells were grown for 1 day with Stem Differentiation Neuronal induction medium (Stem Cell Technologies Catalog #05835), supplemented with the 10 μM ROCK inhibitor, Y27632 dihydrochloride (Tocris Catalog #129830-23-2), to ensure the survival of singly dissociated stem cells. The following day, a confluent monolayer formation was observed and cells were switched to a neuronal induction medium (NIM) made of a 1:1 DMEM/F-12 GlutaMAX (Life technologies, Catalog #10565042) and Neurobasal (Life technologies, Catalog #21103049) with 1x N2 supplement (Life technologies Catalog #17502048), 1x B27 supplement (Life technologies Catalog #17504044), insulin (5µg/ml, Sigma, Catalog #I9278-5ML), L-glutamine (1mM, Life technologies, Catalog #25-030-081), sodium pyruvate (500 μM, Sigma, Catalog #113-24-6), nonessential amino acids solution (100 μM, Life technologies, Catalog #11-140-050), β-mercaptoethanol (100 µM, Sigma, Catalog #M3148-100ML), penicillin-streptomycin (50U/ml, Life technologies, Catalog #15-140-122), Dorsomorphin (1µM, Stemgent, Catalog #04-0024) and SB431542 (10µM Stem cell technologies, Catalog #72232). After neuronal induction was assessed to be successful by the presence of neuronal rosettes (between day 12 to 16), neuronal progenitors were expanded with the addition of recombinant FGF2 (10µM PeproTech Catalog #100-18C) from days 17 to 21. For final differentiation, neurons were plated on laminin (20µg/ml, Thermo-Fisher, Catalog #23017015) and poly-ornithine (Sigma catalog #P4957-50ML) coated plates. Neurons were allowed to mature in neural maintenance media (NMM), containing all the components of NIM with the exception of Dorsomorphin and SB431542. After day 21, neurons were fed every other day with half media changes.

### Acute knockdown of ASH1L in HEK cells and hESC derived human excitatory neurons

shRNA lentiviral constructs targeting ASH1L, containing a puromycin resistance cassette were purchased from Sigma (Supplementary Table S7). Constructs were first tested in HEK cells to assess the knockdown levels, using lipofectamine 2000 (Invitrogen catalog # 11668027) with 1µg of shRNA construct per well of a 6-well plate of HEK cells that were 75% confluent. Puromycin (*In vivogen* catalog # ant-pr-1) selection on HEK cells transfected with shRNAs against ASH1L was started a day after transfection and continued for 3 days with 0.2 µg/ml puromycin concentration. hESCs human cortical neurons were nucleofected with shRNAs constructs against ASH1L using the X-unit of the Amaxa nucleofector unit (Lonza, catalog # AAF1002X). Human neurons between days 30 to 32 of neuronal induction were single cell dissociated using single cell dissociation reagent (stem cell technologies CT catalog # 07174) counted after dissociation, aliquoted into 400,000 cells per aliquot, and centrifuged at 200g for 5 minutes. Cells were resuspended into complete nucleofection solution P3 (Lonza catalog # V4XP-3032) to which a total amount of 1µg of final DNA was added using either 0.8µg of puromycin containing vector or luciferase shRNA in a puromycin resistance vector + 0.2µg of GFP vector, 1µg of GFP vector alone or 0.8µg of shRNA constructs + 0.2µg of GFP vector. Nucleofections were carried out in a final volume of 20µl total using the DC-100 pre-programmed program. Nucleofected cells were allowed to recover for 10 minutes at room temperature after nucleofection and then were resuspended in 80µl of pre-warmed NMM and plated either entirely into 1 well of a 24 well plate for RT-qPCR experiments or plated as 100,000 cells per well in a well of an 8-well chamber slide (Lab-Tek catalog # 177445) for microscopy analysis. After 24 hours nucleofected cells were treated with puromycin at 0.1µg/ml for the first 48hrs, then 0.05µg/ml for an additional 24hrs, and were harvested after 72 hr of the initial Puromycin treatment cells were harvested for either RT-qPCR or fixed for immunocytochemical analysis.

### Western Blotting for ASH1L *in vitro* expression analysis

Protein samples were prepared from cell lysates isolated in ice-cold RIPA buffer [50 mM Tris/HCl (pH 7.4), 150 mM NaCl, 5 mM EDTA, 0.5% (v/v) NP40 (Nonidet P40), containing protease inhibitor and phosphatase inhibitor (Pierce Catalog #P5726-1ML). Proteins were quantified by the BCA protein assay kit (Pierce Chemical Co. catalog # 23227). A total of 20 to 40µg of protein were loaded per lane and separated by 4-12% Bolt-PAGE gels (Invitrogen catalog # NW04122BOX) and transferred to nitrocellulose membrane in Towbin buffer that contained 25mM Tris, 192mM glycine, and 20% (v/v) Methanol (pH8.3) for 1.5 hour on ice at constant current (450mA). Membranes were probed with the following antibodies: anti-ASH1L antibody (Bethyl laboratory catalog # A301-748A), anti-PAX6 (Abcam catalog # ab5790), anti-α-TUBULIN antibody (sigma catalog # T9026), anti-NANOG (Abcam catalog # ab21624) and anti-MAP2 (Abcam Catalog #ab5392). Species-specific secondary antibodies coupled to HRP molecules were used at 1:10,000 (mouse) or 1:15,000 (rabbit) dilutions (Jackson immune laboratories). The bands were visualized by chemiluminescence detection system (Thermofisher sci. catalog # 34577) and analyzed using Fiji/ImageJ (Schindelin et al., 2012; Schneider et al., 2012).

### RT-qPCR analysis of gene expression

Total RNA was harvested from hESCs and hESCs-derived neural cultures using a total RNA mini-solation kit (Qiagen catalog #74104). RNA was reverse transcribed to cDNA using the Superscript reverse transcription kit (Invitrogen catalog #18080-051), following manufacturer’s recommendations. Gene expression levels were determined using SYBER green with IDT designed primers (Supplementary Table S8) for detection of pluripotency markers (*OCT4*), neuronal progenitor marker (*PAX6*) or neuronal markers (*MAP2*) or neurotrophin receptors (*NTRK1, NTRK2, NTRK3* and *P75*). SYBR green reactions were performed following the manufacturer’s protocols in a CFX96 Touch Real Time PCR Detection System (BioRad) using 4ng of total cDNA, and 1µM of SYBR Green intercalator primers (Thermofisher catalog #4367659). qPCR reactions were run using the following parameters: 95°C for 10 minutes, followed by 40 cycles of 95°C for 15 seconds and 60°C for 1 minute. Single amplicons of 100 to 200 bp were produced with no amplification from genomic DNA. For all reactions three technical replicates (replicate wells of qPCR) were used. The BioRad software determined Ct values automatically. We analyzed the results using the ΔΔCt method, normalizing the data to the reference gene GAPDH, this data was then further normalized to control GFP. p-values were calculated based on the normalized expression levels (ΔΔCt) using paired t-tests.

### ddPCR analysis of TrkB isoforms

To conduct ddPCR analysis, primers were custom designed for specific TrkB isoforms (Supplementary Table S9). Primers and probes were purchased from IDT technologies. 0.5 to 5ng of cDNA were used for each experimental condition in 20µl reactions containing primers, probes and ddPCR mix were used to generate individual droplets using a QX200™ Droplet Generator (Bio-Rad). Reactions were then subjected to PCR amplification using the following conditions: 95°C for 5 min, followed by 40 cycles of 95°C for 30 seconds and 60°C for 1 min (ramp rate was 2°C/s), then 4°C for 5 minutes, 90°C for 5 minutes for signal stabilization using a QX200 Droplet Digital PCR (ddPCR™) System – Bio-Rad. ddPCR data was analyzed by normalizing to GAPDH and then normalizing to control GFP to account for transfection efficiency.

### Immunocytochemistry of hESCs derived forebrain cortical neurons

Nucleofected human neurons were immunostained as previously described (Ouyang et al., 2013). Briefly, neurons were fixed in 4% paraformaldehyde for 15 to 30 minutes at room temperature, rinsed three times with 1xPBS, and cell membranes were permeabilized with 2% triton in 1XPBS for 10 minutes and then rinsed as above. Cells were blocked with 10% goat serum diluted in 1xPBS for 1 hr at room temperature, and incubated with primary antibodies diluted in 2% goat serum, 0.25% 1xTriton and 1xPBS for either 3 hrs. at room temperature or overnight at 4°C. Primary antibody was removed by 3 washes of 5 minutes each with 1xPBS. Cells were then incubated with Alexa secondary antibodies (Invitrogen) diluted at 1:800 in 1xPBS. Neurite analysis was conducted by immunostaining with β-III-Tubulin (Millipore, catalog #AB9354 at 1:500) or MAP2 (Abcam catalog #ab5392 at 1:500). Activation of the BDNF-TrkB downstream signaling was assessed by analysis of CREB phosphorylation using an anti-Phospho-CREB antibody (Cell signaling, catalog #9198S at 1:500).

### EZH2 and BDNF treatment of ASH1L knockdown neurons

To determine the capacity of BDNF or EZH2 inhibitors to rescue the ASH1L associated neuronal phenotypes, we treated nucleofected neurons 48hrs after the initial puromycin treatment with either BDNF (Preprotec, Catalog #450-02 at 10ng/ml) or EZH2 inhibitor EI1 (Sigma-Aldrich /Calbiochem Catalog #5005610001 at 1.25 µM and 2.5µM). Treatments were carried out for 72hrs in BDNF and both 72hrs and 120hrs with EZH2 inhibitor. After the initial treatment neurons were fed with half media changes containing the final concentration of either BDNF or EZH2 inhibitor.

### Phalloidin Staining

In order to analyze changes in growth cone number and size, after 72hrs of puromycin treatment nucleofected neurons were stained with phalloidin. Briefly, neurons were fixed for 30min with 4% paraformaldehyde and then processed as described in the immunofluorescence methods section except that after the secondary antibody incubation they were incubated in Alexa-568 Phalloidin (Invitrogen, catalog #A12381, diluted at 1:400) for 30 minutes and then rinsed with 1xPBS three times for 30 seconds each time. Cells were then mounted using Anti-Fade GOLD-DAPI (Thermo-Fischer, catalog #P36931).

### Image Acquisition

For neuronal arborization and growth cone studies images were taken using a Leica DMI3000 fluorescence microscope or with a Leica DM6000 widefield fluorescence microscope. For analysis of phospho-CREB signal, cells were imaged with conventional widefield microscopy with deconvolution. In GFP-expressing neurons, super resolution images of the nucleus were taken using structured illumination microscopy (DeltavisionOMX-SR). This technique was used in order to ascertain the distribution of Phospho-CREB in the nucleus. To determine the levels of phospho-CREB, we took images using a widefield fluorescence microscopy at 40 and 63X magnifications.

### Image Analysis

To analyze neuronal arborization parameters we utilized Neurolucida software and traced GFP stained neurons after confirming that they were positive for β-III-Tubulin. Observers carrying out the tracing were blinded to experimental conditions. Neurons were traced using the Neurolucida 360 program after adjusting for the pixel to µm ratio for both the x and y parameters. The cell bodies were detected using the “detect all soma” button on the cell body panel. The dendrites were then traced by changing the controls to “user guided kernels”, following each dendrite starting at the cell body. To analyze traced neurons, the Neurolucida Explorer program was used.

To quantify the total number of growth cones and the size of the growth cone areas, we analyzed images using ImageJ by using the threshold function and drawing regions of interest around phalloidin stained growth cone-like areas. The area, perimeter, and integrated density of each region of interest was then measured.

To quantify the total content of phospho-CREB translocated to the nucleus, images were also analyzed using ImageJ. Images for DAPI and phospho-CREB were changed to 16-bit. Regions of interest (ROI) were drawn around each nucleus in the DAPI channel and then added to the ROI manager and loaded onto the phospho-CREB channel and measured. A background area was also measured for each phospho-CREB image. Mean fluorescent intensity (MFI) was calculated by using the following formula: ***MFI = Integrated density of phospho-CREB – (phospho-CREB area*mean grey value background)***

### Statistical Analysis

Unless otherwise specified, all data was analyzed using GraphPad statistical analysis software and multiple groups were either analyzed using one-way ANOVA with multiple comparisons without correction when compared to a single control or using a two-way ANOVA with multiple comparisons without correction for multiple comparisons when multiple treatments or conditions were compared or using paired or unpaired t-tests.

### Identification of epigenetic mark content in neurotrophin receptors

Epigenetics data was compiled from the NIH Roadmap Epigenomics Mapping Consortium (https://egg2.wustl.edu/roadmap/web_portal/processed_data.html). 127 consolidated epigenomes were analyzed to create the datasets. Genome-wide signal coverage tracks for H9 derived neuron cultured cells were visualized for all the neurotrophin receptors using -log10 (p-value) signal tracks. The receptors of interest included NTRK1, NTRK2, NTRK3, and p75. For these genes, the histone marks H3K27me3, H3K36me3, and H3K4me3 were available in the H9 derived neurons.

## Supporting information

Cheon_etal_2020BioRXV_SupplementalTables

## ACKNOWLEDGEMENTS

We thank Jeffrey Twiss for critically reading the manuscript. We thank Jeffrey Twiss and Amar Kar for advice on how to quantify the phospho-CREB nuclear signal. This work was supported by the Center of Biomedical Excellence Dietary Supplements and Inflammation-NIGMS P20GM103641, and SC INBRE NIGMS P20GM103499 Pilot Award to S.B.L. Research reported in this publication was supported in part by the SC EPSCoR/IDeA Program under award number 18-SR04 to S.B.L. The views, perspective, and content do not necessarily represent the official views of the SC EPSCoR/IDeA Program. JSL is supported by 1R01NS104428-01A1.

## AUTHOR CONTRIBUTIONS

SBL conceived, designed and supervised the study. SHC conducted the majority of the experiments. SHC and SBL conducted the majority of the analysis. SBL and AMC conducted additional EZH2 rescue, phalloidin, and phospho-CREB signaling experiments and, FDR contributed to some of these experiments. AMB conducted the Allen brain atlas and ASH1L correlation analysis. AMB and SHC conducted the majority of the morphogenesis analysis, and MHC contributed to this analysis, too. AMC, FDR, MM, and AJS conducted imaging and analysis of phospho-CREB, phalloidin and additional EZH2 experiments. JMC conducted the analysis of the epigenetic marks in ENCODE. CMP & JSL conducted the imaging on the Delta vision OMX microscope. EC & PSG provided some of the neurons used in some of the experiments. SBL wrote the manuscript and put together all the final figures and tables. All authors contributed to editing the manuscript and interpretation of the results.

## SUPPLEMENTARY MATERIAL

**Supplementary Figure S1.**
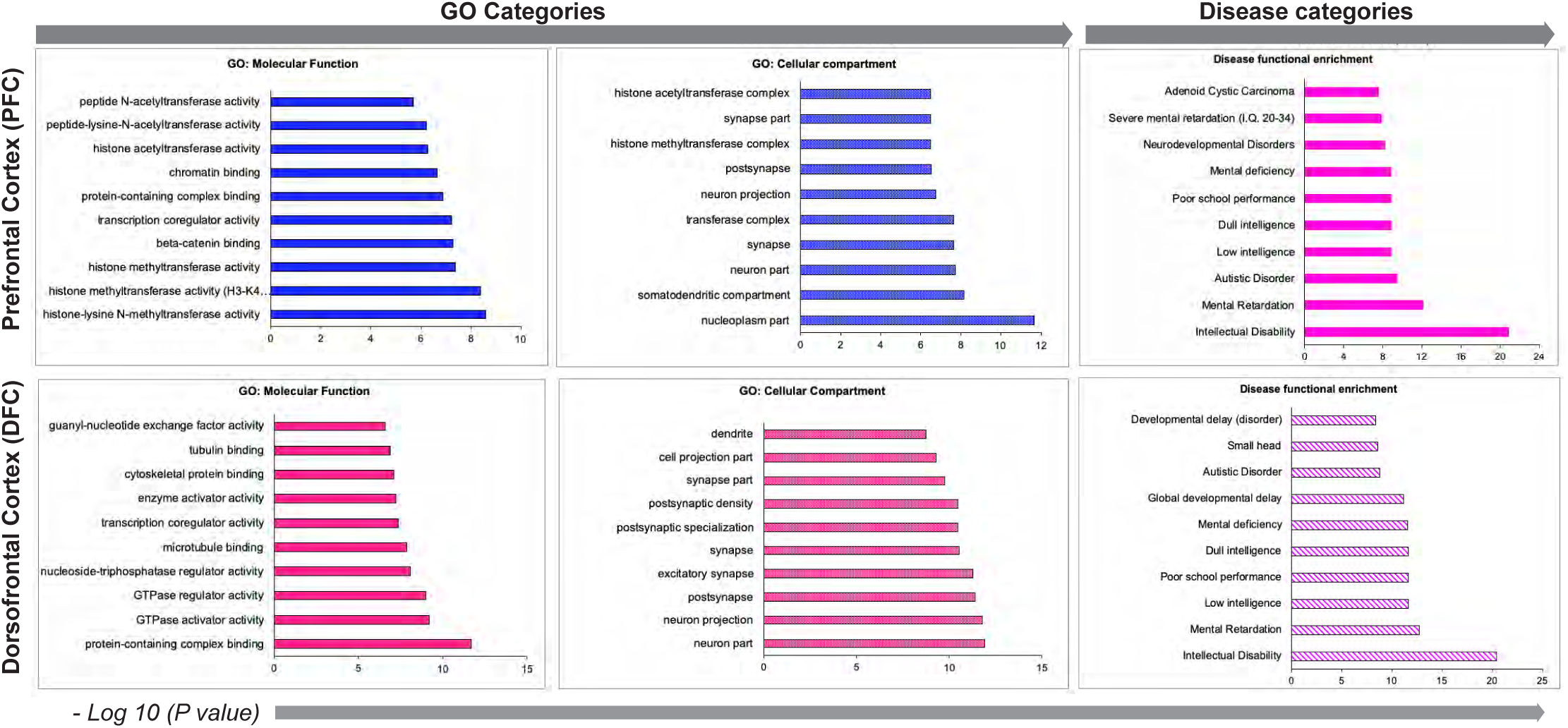
*(related to figure 1 in main text).* Pathway analysis of Top-500 correlated genes with *ASH1L.* The top 10 pathways by most significant adjusted p-value for GO categories for Molecular function and Cellular compartment and for Disease categories are shown for the prefrontal Cortex (PFC) or Dorsofrontal cortex (DFC). p-values are shown as the negative Log10 on the X-axis. GO and disease categories are in the Y-axis.

**Supplementary Figure S2.**
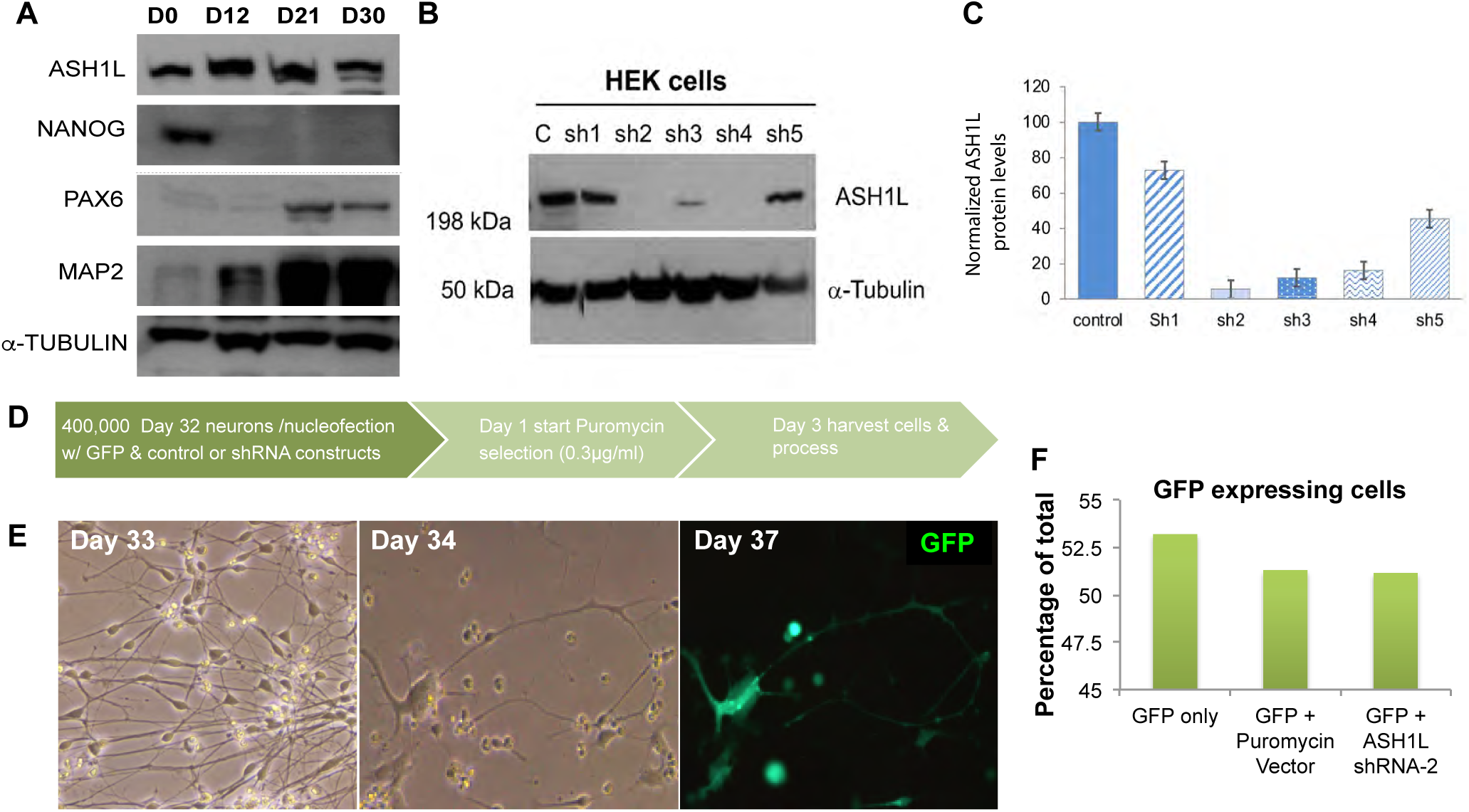
*(related to figures 1 and 2 in main text)*. ASH1L protein expression in vitro, knockdown of ASH1L in HEK cells and nucleofection of ASH1L in cortical human neurons. (**A**) Representative western blot shows ASH1L protein expression at different *in vitro* developmental stages marked by expression of NANOG (stem cell), PAX6 (neuronal progenitors), and MAP2 (neurons) with α-Tubulin as control. (**B**) Representative western blot shows ASH1L protein expression after knockdown with 5 different ASH1L targeting shRNA constructs (sh1, sh2, sh3, sh4 and sh5) control lane (C) is untransfected control, top blot shows ASH1L band above the 198 Kilo Dalton (kDa) marker and bottom image shows the α-Tubulin control band around the 50 kDa molecular weight marker. (**C**) ASH1L protein expression levels are shown in HEK cells after transfection and puromycin selection as a percentage of control. (**D**) Experimental design shows nucleofection of day 29 to 32 cortical neurons with GFP and shRNA constructs followed by puromycin selection with 0.1 to 0.3µg/ml concentration and harvesting of neurons 72hrs after initial puromycin treatment. (**E**) Images of nucleofected neurons after puromycin selection on days 33 to 34. (**F**) Analysis of nucleofection efficiency by quantification of GFP expressing cells as a ratio of total cells.

**Supplementary Figure S3.**
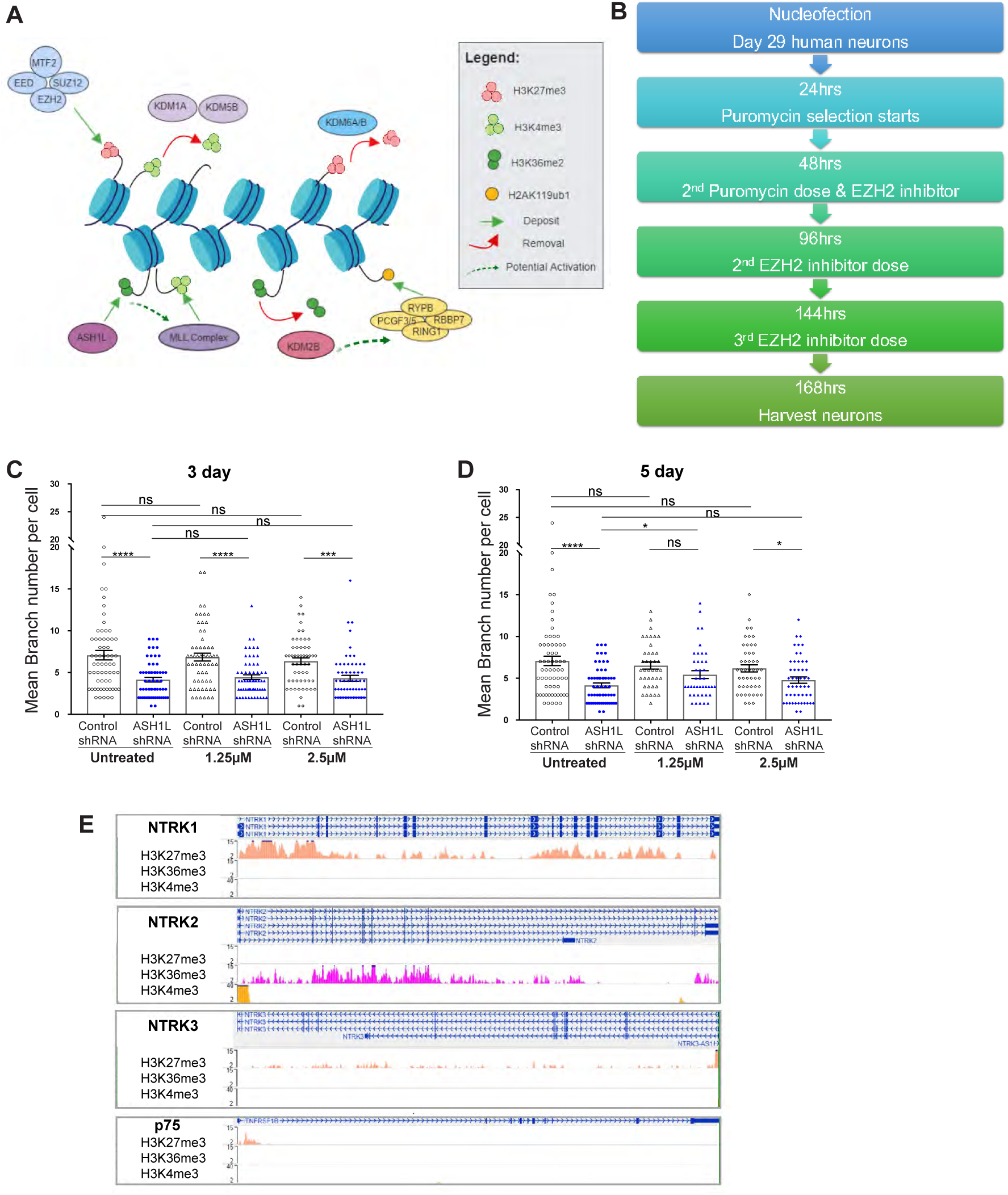
*(related to figure 3 in main text).* Epigenetic regulation of neurotrophin receptors and description of TrkB isoforms analyzed in this study. (**A**) Diagram representing the counteracting activity of the Polycomb and Trithorax genes and their epigenetic marks in chromatin. Red circles represent repressive marks H3K27me3, light green circles represent activating marks H3K4me3 and dark green circles represent activating marks H3K36me2. PRC2 complex is shown in light blue, PRC1 complex is shown in yellow and members of the Trithorax group are shown in different purple shades. (**B**) Experimental design for EZH2 inhibition in neurons with ASH1L knockdown. (**C**) Mean branch number per cell was analyzed in cortical neurons under control or ASH1L knockdown conditions that were either untreated or treated for 3 days with 1.25µM or 2.5µM EZH2 inhibitor. All measurements per cell are shown for each condition in combination with the mean ± SEM. Control-shRNA (open black circles) = 7.08 ± 0.56 *vs* ASH1L-shRNA (solid blue circles) = 4.15 ± 0.28; Control-shRNA+1.25µM EI1 (open black triangles) = 6.85 ± 0.48 *vs* ASH1L-shRNA +1.25µM EI1 (solid blue triangles) = 4.45 ± 0.29; Control-shRNA + 2.5µM EI1 (open black diamonds) = 6.36 ± 0.40, n = 48 cells *vs* ASH1L-shRNA-2 + 2.5µM EI1 (solid blue diamonds) = 4.31 ± 0.35. ****P < 0.0001, ***P < 0.0007, not significant = ns. (**D**) Mean branch number per cell was analyzed in cortical neurons under control or ASH1L knockdown conditions that were either untreated or treated for 5 days with 1.25µM or 2.5µM EZH2 inhibitor. All measurements per cell are shown for each condition in combination with the mean ± SEM. Control-shRNA (open black circles) = 7.08 ± 0.56 *vs* ASH1L-shRNA (solid blue circles) = 4.15 ± 0.28; Control-shRNA+1.25µM EI1 (open black triangles) = 6.52 ± 0.43, *Vs.* ASH1L-shRNA +1.25µM EI1 (solid blue triangles) = 5.44 ± 0.47, *p = 0.123*; Control-shRNA + 2.5µM EI1 (open black diamonds) = 6.19 ± 0.42 *vs* ASH1L-shRNA+ 2.5µM EI1 (solid blue diamonds) = 4.78 ± 0.38. ****P < 0.0001, *P < 0.05. (**E**) Analysis of neurotrophin receptors encoding genes NTRK1, NTRK2, NTRK3 and p75 in NIH Roadmap Epigenomics Mapping Consortium database. Analysis was done for H9 cortical excitatory neurons, which are female neurons. Multiple isoforms are shown for each gene. H3K27me3 peaks are shown in light orange, H3K36me3 peaks are shown in hot pink, and H3K4me3 peaks are shown in yellow.

**Supplementary Figure S4.**
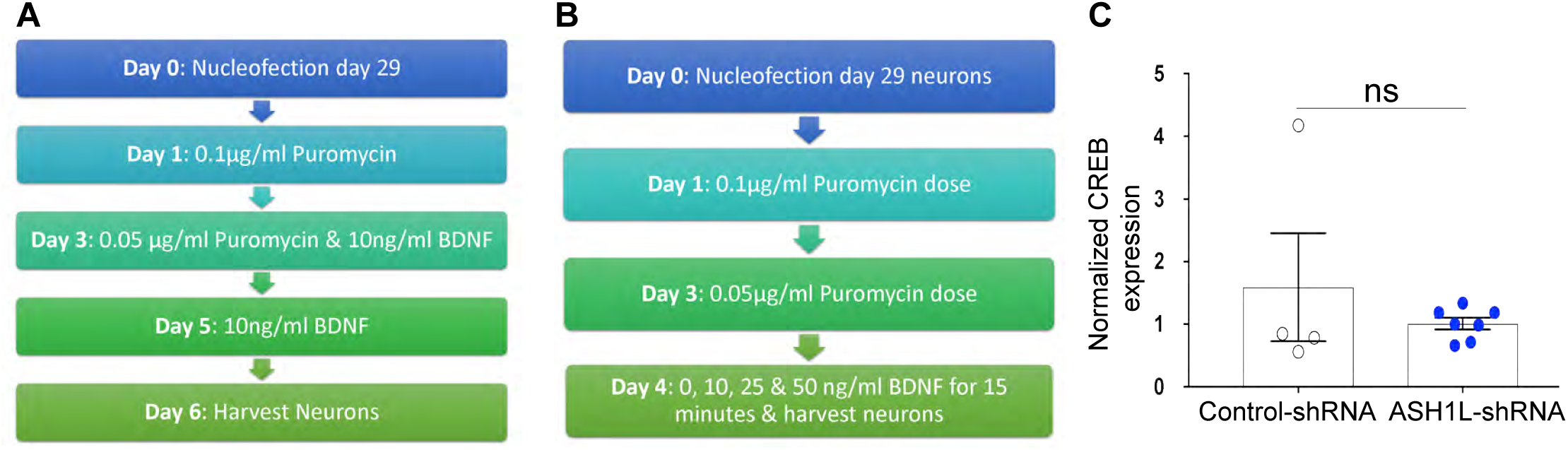
*(related to figure 4 in main text).* Extended and Acute BDNF treatment in ASH1L knockdown neurons, Phospho-CREB activation and CREB expression analysis. **(A)** Diagram depicting the BDNF long term rescue experiment in control-shRNA and ASH1L-shRNA nucleofected neurons. **(B)** Diagram depicting the acute BDNF experiment in control-shRNA and ASH1L-shRNA nucleofected neurons at varying doses of BDNF. **(C)** Knockdown of ASH1L in male cortical neurons does not alter the mRNA levels of CREB1. RT-qPCR analysis of gene expression shown as normalized data to reference gene GAPDH that was then normalized to control GFP is shown as ΔΔCt values for control-shRNA (open black circles, n=4 independent experiments) and ASH1L shRNA (blue solid circles, n=7 independent experiments) for CREB1. All statistical analysis was carried out using an unpaired t-test, ns= 0.39. Mean values are shown by the bars and error bars are SEM.

## SUPPLEMENTARY TABLES

**Supplementary Table-1.**
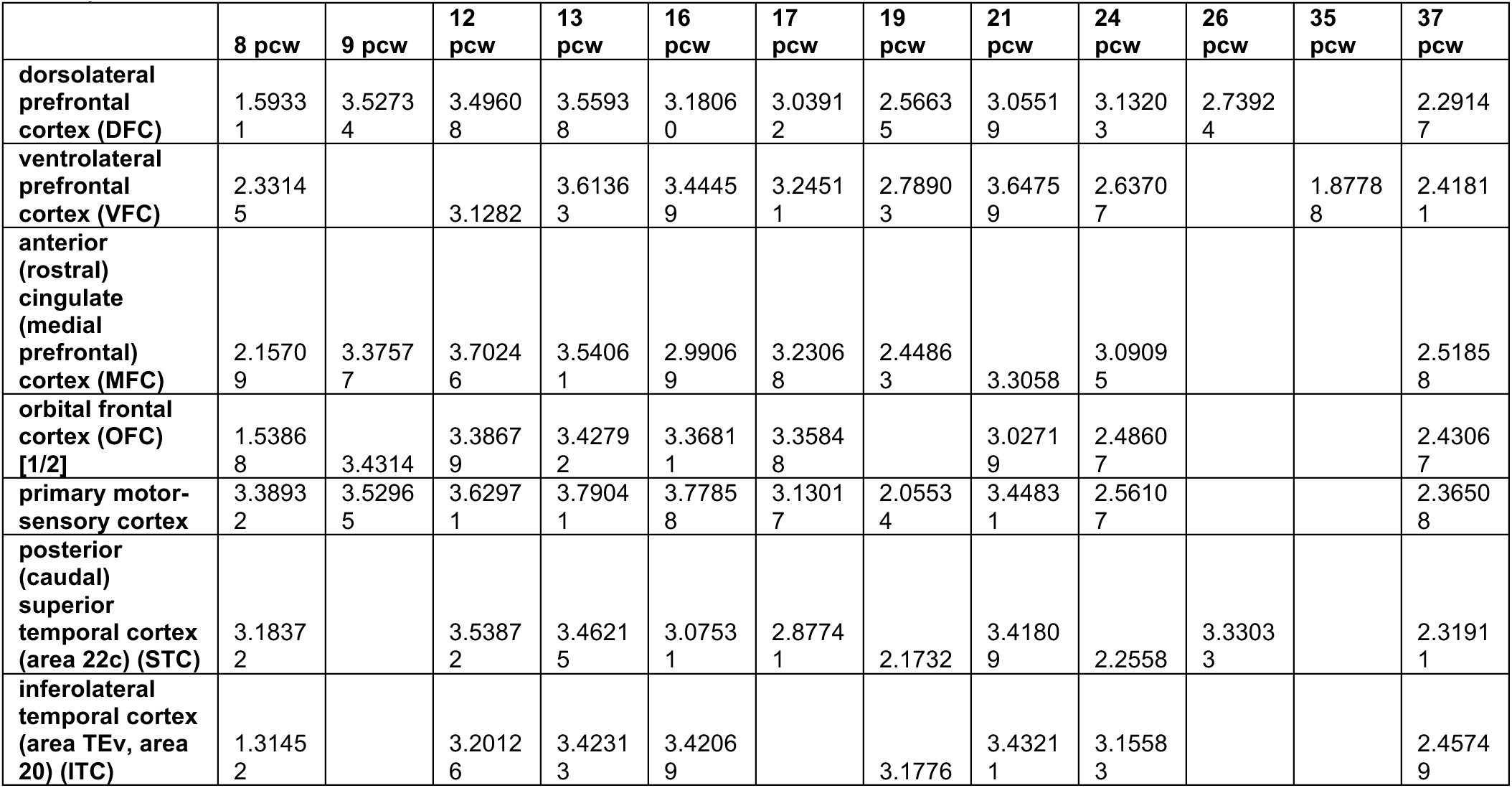
***(related to figure 1 in main text).* Allen Brain Atlas heat map average expression data per post-conception week (pcw).** Cortical areas analyzed for ASH1L expression from 8 to 37 post-conception weeks fetal brain.

**Supplementary Table-2 *(related to figure 1 in main text).* Allen Brain Atlas raw expression data per donor in all areas for each pcw.** All raw data used for all areas used in the analysis for all donors used (excel file format).

**Supplementary Table-3.**
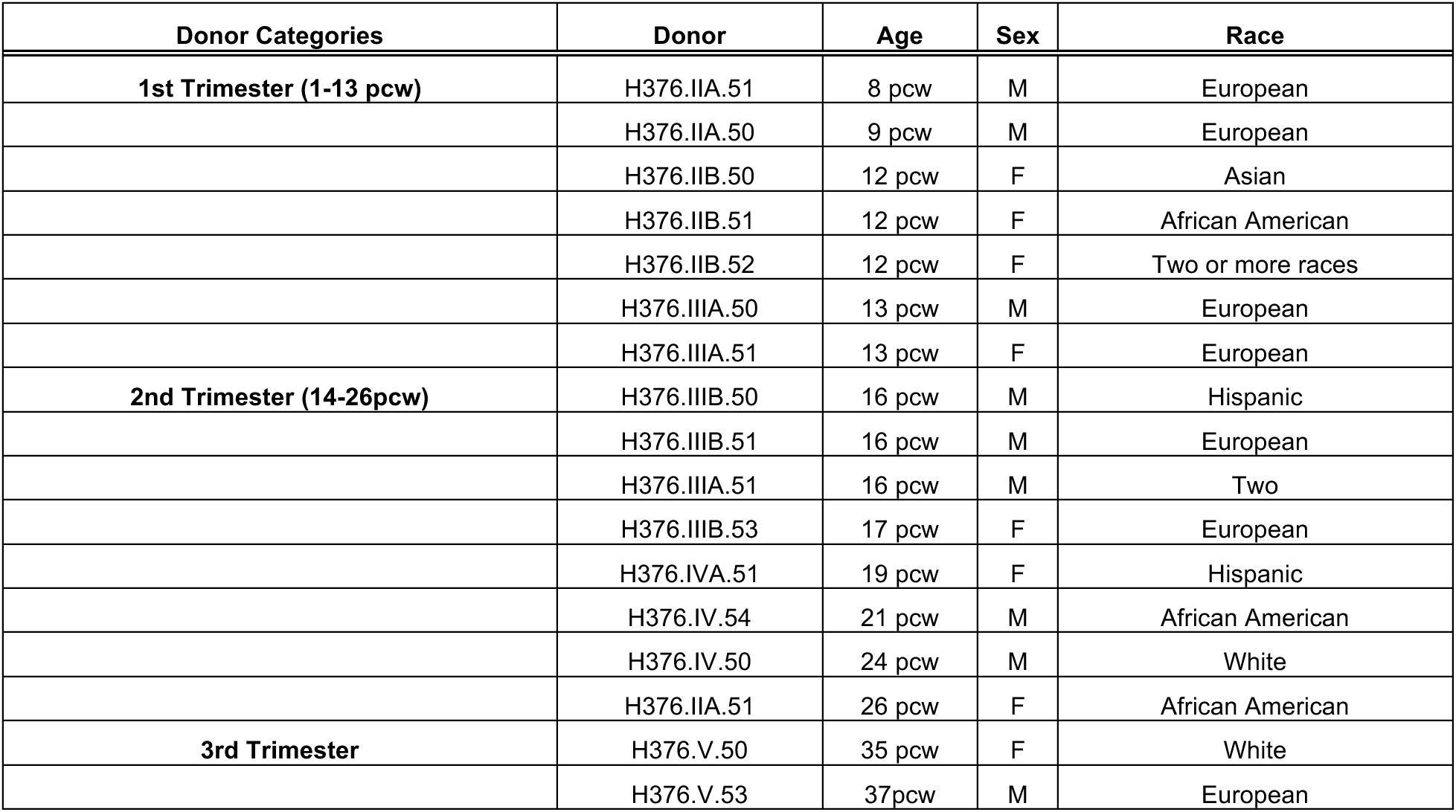
***(related to figure 1 in main text).* Donor information from Allen brain Atlas.** Information per donor showing age in post-conception weeks (pcw), sex and race.

**Supplementary Table-4 *(related to figure 1 in main text).* Top 500 co-expressed genes with ASH1L in the prefrontal cortex and the dorsal frontal cortex.** Top co-expressed genes are shown by gene symbol, gene name and R correlation co-expression coefficient for the prefrontal cortex (PFC) and the dorsal frontal cortex (DFC) (excel file format).

**Supplementary Table-5 *(related to figure 1 in main text). Toppfun analysis of* Top 500 co-expressed genes with ASH1L in the Prefrontal cortex (PFC).** Pathway analysis showing GO categories for molecular function, biological process, and cellular compartment. In addition, association to human phenotype, pathways and disease are also shown for the Prefrontal cortex gene cohort (excel file format).

**Supplementary Table-6 *(related to figure 1 in main text). Toppfun analysis of* Top 500 co-expressed genes with ASH1L in the Dorsal Frontal Cortex (DFC).** Pathway analysis showing GO categories for molecular function, biological process, and cellular compartment. In addition, association to human phenotype, pathways and disease are also shown for the Dorsal frontal cortex gene cohort (excel file format).

**Supplementary Table-7.**
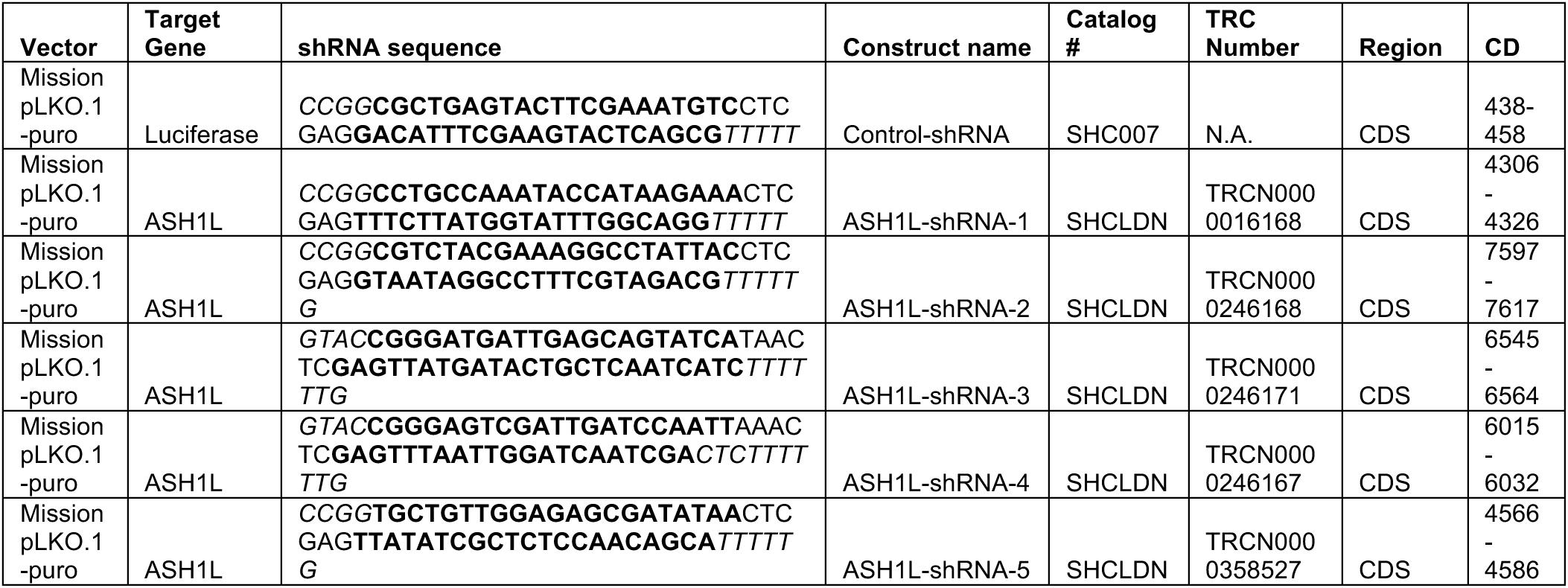
***(related to figures 1,2,3 and 4 in main text). shRNA constructs used in this study*.** Pathway analysis showing GO categories for molecular function, biological process, and cellular compartment. In addition, association to human phenotype, pathways and disease are also shown for the Dorsal frontal cortex gene cohort.

**Supplementary Table-8.**
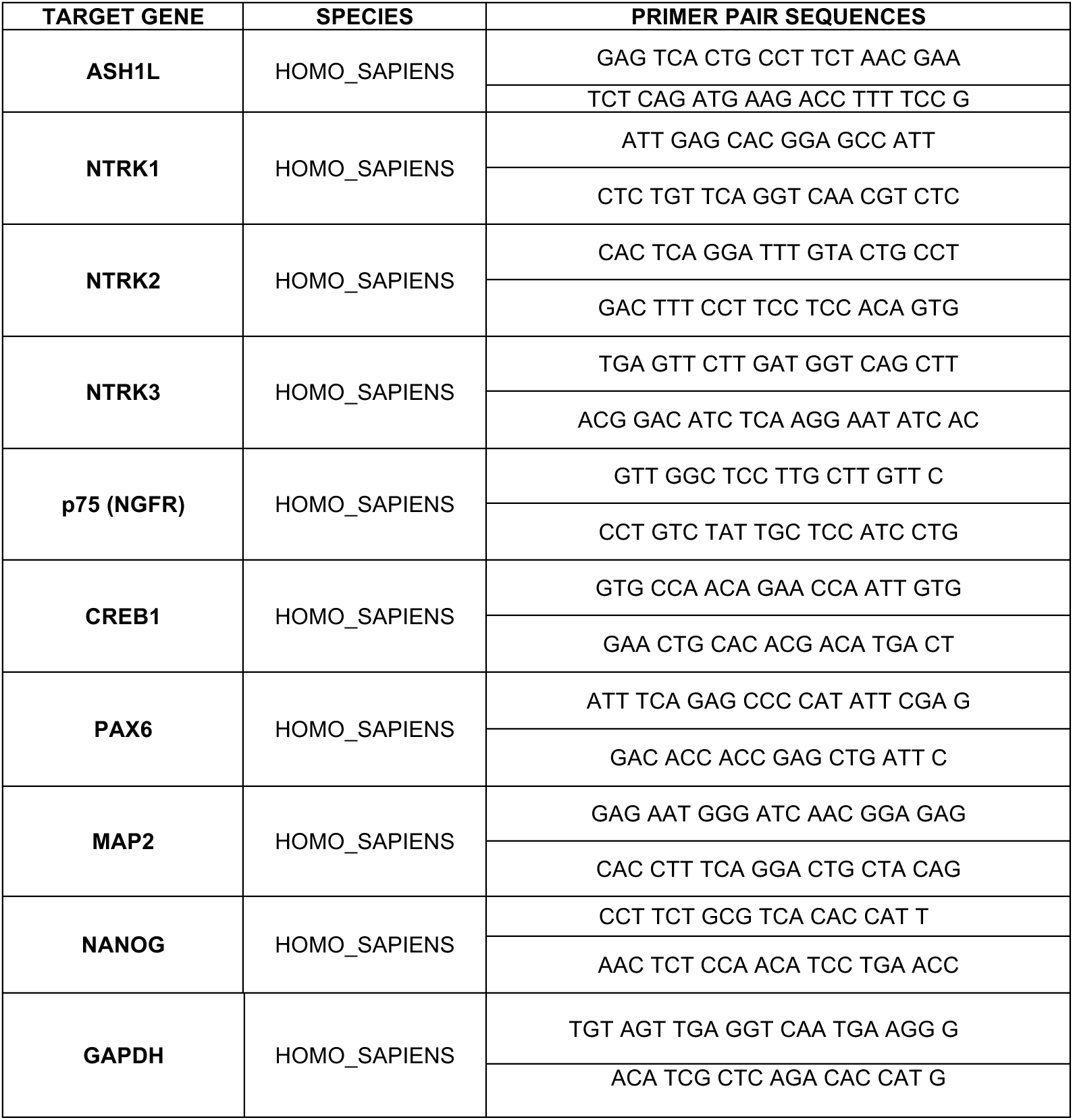
***(related to figures 1 and 3 in main text and supplementary figure S4).* Quantitative PCR primers used for gene expression analysis.** All primer pair sequences used for qPCR experiments are described in this table.

**Supplementary Table-9.**
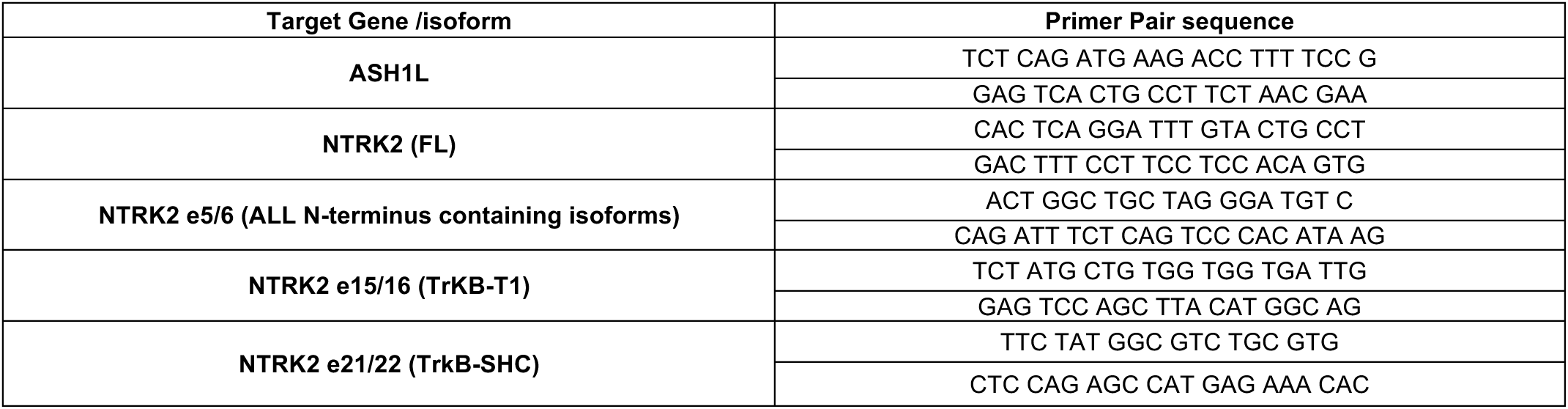
***(related to figure 3 in main text).* NTRK2 isoform primers for digital droplet PCR (dd-PCR).** All primer pair sequences used for dd-PCR experiments to analyze the different NTRK2 (TrkB) isoforms are described in this table.

